# Protein Language Models and Structure-Based Machine Learning for Prediction of Allosteric Binding Sites in Protein Kinases: An Explainable AI Framework Grounded in Energy Landscape-Encoded Frustration

**DOI:** 10.64898/2026.01.05.697819

**Authors:** Kamila Riedlová, Vít Škrhák, Will Gatlin, Max Ludwick, Lucas Turano, Marian Novotný, David Hoksza, Gennady M. Verkhivker

**Affiliations:** Department of Software Engineering, Faculty of Mathematics and Physics, Charles University, Prague, Czech Republic; Keck Center for Science and Engineering, Graduate Program in Computational and Data Sciences, Schmid College of Science and Technology, Chapman University, Orange, CA 92866, United States of America; Department of Biomedical and Pharmaceutical Sciences, Chapman University School of Pharmacy, Irvine, CA 92618, United States of America; Department of Pharmacology, Skaggs School of Pharmacy and Pharmaceutical Sciences, University of California San Diego, 9500 Gilman Drive, La Jolla, CA 92093, United States of America; Department of Cell Biology, Faculty of Science, Charles University, Prague, Czech Republic

**Author notes:** Correspondence; (G.V).

## Abstract

Reliable identification of allosteric binding sites remains a major bottleneck in structure-based drug discovery, particularly in protein kinase families where such sites are often structurally cryptic, evolutionarily non-conserved, and sparsely populated. In this work, we present a systematic analysis of binding site prediction across a rigorously curated dataset of human kinase–ligand complexes, encompassing 453 kinases and spanning five inhibitor classes: Type I, Type I.5, and Type II (orthosteric ATP-competitive) and Type III/IV (non-ATP allosteric) modulators. We employed the pretrained protein language model (PLM) ESM2-650M model that was fine-tuned for prediction of protein-ligand binding sites by replacing the original masked language modeling head with a token-level classification head that acts as a projection layer that maps the high-dimensional latent representation of each residue to a scalar probability score for a given protein residue to be part of the binding site. We employed this fine-tuned sequence-based PLM and structure-based detection approach P2Rank for identification of orthosteric and allosteric binding sites in protein kinases. Our analysis reveals a stark performance divergence: while both methods achieve high precision-recall (AUPR = 0.64-0.76) on orthosteric sites, PLM performance collapses on allosteric sites (AUPR = 0.06), despite retaining moderate ranking ability (AUROC = 0.70). This deficit persists even after strict control for sequence similarity, structural redundancy, and extreme class imbalance (allosteric residues constitute <3% of the kinase domain). To mechanistically interpret this discrepancy, we integrate large-scale local frustration analysis, a physics-based framework derived from energy landscape theory that quantifies the energetic stability of residue-residue interactions under mutational and conformational perturbations. We find that, although the global frustration landscape is conserved across kinase states dominated by neutral frustration (55-75% of residues), local binding sites exhibit fundamentally distinct mutational constraints. Orthosteric pockets are enriched in minimally frustrated residues, whereas allosteric sites are characterized by neutral mutational frustration, indicating evolutionary permissiveness and sequence degeneracy. This study reframes the performance of AI approaches in predicting protein binding sites as a reflection of functional design that can be rationalized through lens of the landscape-encoded protein frustration as an explainable AI framework.

## 1. Introduction

The discovery of druggable binding sites in proteins—particularly those that are cryptic, transient, or allosteric—remains one of the most persistent bottlenecks in structure-based drug design.^1–5^

Over the past decade, computational strategies for predicting binding sites and allosteric pockets have evolved into a multi-faceted toolkit, categorized into geometric, energetic, evolution-based, and generative machine learning (ML) approaches. While traditional template-free methods like Fpocket^6^, SiteHound^7^, CurPocket^8^ and MetaPocket 2.0^9^ remain widely used for their reliance on surface geometry and probe-based energetics, the specific challenge of identifying allosteric sites—which are often distal to the active site—has necessitated more specialized, dynamics-aware models. With the steady increase in the size, quality and rigorous annotations of biological databases, there has been the growing interest in machine learning (ML) methods for protein–ligand binding site prediction. A template-free P2Rank approach^10,11^ is among best performing ML tools for prediction of ligand binding sites^4^ and is based on prediction of ligandability of local chemical neighborhoods that are centered on points placed on the solvent accessible surface of a protein. P2Rank employs random forest method trained on known protein–ligand complexes to score the “ligandability” of local surface neighborhoods, offering generalization across protein families without requiring templates.^10,11^ The emergence of deep learning has shifted the focus toward structure-based algorithms that can extract binding site features from raw 3D data. A notable example is ISMBLab-LIG, which predicts ligand binding sites by transforming protein surfaces into 3D probability density maps of interacting atoms. By reconstructing these maps on the query surface, the model remains robust against the local conformational flexibility often seen in tentative binding pockets.^12^ Recent advancements have increasingly utilized 3D voxelization—the process of discretizing the protein structure into a grid—to leverage the power of computer vision architectures. DeepSite pioneered this approach by using 3D Deep Convolutional Neural Networks (DCNNs), significantly outperforming traditional structure-based algorithms in large-scale benchmarks involving over 7,000 proteins from the scPDB database.^13^ Building on this voxel-based foundation, subsequent models have adopted specialized architectural frameworks to refine detection. FRSite utilizes a Faster R-CNN (Region-based Convolutional Neural Network) architecture, treating the identification of binding sites as a 3D object detection task.^14^ In contrast, Kalasanty employs a U-Net segmentation architecture, which is widely recognized in medical imaging for its ability to produce high-resolution, residue-level classifications of binding volumes.^15^ BiteNet (Binding site neural Network), a deep learning approach designed for the rapid, spatiotemporal identification of protein pockets whereby by treating protein conformations as 3D images and conformational ensembles as 3D videos, the method leverages computer vision principles to detect binding sites as “objects” within a dynamic structural landscape.^16^ Unlike static methods, BiteNet excels at capturing the structural fluctuations of soluble and transmembrane proteins, making it particularly effective for identifying allosteric and cryptic sites across complex targets, including conformation-specific allosteric site in the EGFR and an oligomer-specific site in the trimeric P2X3 receptor.^16^ When validated on the HOLO4K benchmark, BiteNet demonstrated accuracy that was comparable to DeepSite and P2Rank approaches.^17^ Collectively, these models represented a transition from manual feature engineering toward the end-to-end spatial learning of binding pocket determinants. Binding sites that exhibit significant conformational changes are often referred to as cryptic binding sites (CBS). The evaluation of structure-based method P2Rank^10,11^ on the recently developed CryptoBench benchmark^18^ demonstrated a reduction in prediction performance on apo protein conformations in the absence of dynamic signals embedded in ligand-induced changes leading to novel pockets.

The accurate prediction of protein binding sites, including canonical active sites, cryptic functional pockets, and allosteric binding regions—is a cornerstone of structure-based drug discovery. Traditional computational methods (e.g., geometric pocket detection, molecular dynamics simulations, evolutionary conservation analysis) have long faced limitations in scalability, sensitivity to conformational dynamics, and generalizability across protein families. In recent years, the advent of deep learning, particularly large language models (LLMs) and protein language models (PLMs), has catalyzed a paradigm shift in this domain.^19,20^ Notably, PLMs have shown great promise in binding site prediction.^21–25^ Graph Neural Network (GNN)-based method Graph Attention Site Prediction (GrASP) performs a rotationally invariant featurization of solvent-accessible atoms. In the framework of GrASP approach, the authors have created a new publicly available version of the sc-PDB database containing 26,196 binding sites across 16,889 protein structures.^22^

We previously explored the potential of PLMs to predict protein–ligand binding residues, showing superior performance compared to state-of-the-art methods on multiple datasets.^23^ Our latest study explored a hybrid approach that combined PLMs and Graph Neural Networks (GNNs) by constructing a residue-level Graph Attention Network (GAT) model based on the protein structure that uses pre-trained PLM embeddings as node features.^24^ By exploiting a benchmark dataset over a range of ligands and ligand types, we have found that the incorporation of structural information through the GNN improves protein–ligand binding site prediction while for more complex PLMs the relative impact of the structure information represented by the GNN architecture diminishes.^24^ The PLM fine-tuning strategies go beyond static embeddings extracted from the final pretrained PLM, and show improved performance by updating model parameters during training^25^ or through employment of parameter-efficient fine-tuning (PEFT).^26,27^ These PLM finetuning approaches focused on the transformer layers of PLMs and add a small number of new parameters which are tuned, leaving the original model parameters untouched. By requiring significantly fewer resources PLM finetuning strategy can achieve superb performance with smaller computational resources. Our recent studies explored the PLMs with a range of finetuning strategies for predicting cryptic binding sites, evaluated against the baseline to assess the improvement of prediction performance. By incorporating multi-task learning, the results showed sequence-based models can be significantly enhanced through carefully selected finetuning strategies and data sources.^28,29^

As sequence models lack explicit spatial awareness, several effective recent approaches fuse PLM embeddings with 3D structural graphs. EquiBind approach^30^ and related approach DiffSBDD^31^, use equivariant graph neural networks (GNNs) to predict ligand binding poses and pocket locations directly from apo protein structures. Despite the diversity of existing pocket detection tools, identifying allosteric and cryptic binding sites—which often exist only as short-lived intermediate states—remains a formidable challenge. PocketMiner, a deep learning method specifically engineered to identify cryptic pockets that escape detection by traditional crystallography. Unlike previous models, PocketMiner utilizes extensive molecular dynamics (MD) simulations to assess whether individual residues can rearrange their orientation during thermal fluctuations to facilitate pocket opening.^32^

However, PocketMiner showed significantly worse performance compared to simple transfer-learning of the PLM on the CryptoBench benchmark^18^. Among family of generative approaches, PocketGen produces residue sequence and atomic structure of the protein regions in which ligand interactions occur by using a graph transformer for structural encoding and a sequence refinement PLM module.^33^ The graph transformer captures binding interactions while for sequence refinement a structural adapter is integrated into the PLM ensuring alignment between structure-based and sequence-based predictions, and enabling generation of plausible protein pockets with enhanced binding affinity and structural validity.^33^

However, despite rapid advances in AI-driven approaches and PLM strategies, reliable detection, and characterization of cryptic and allosteric pockets across diverse conformational states remains a formidable challenge. This difficulty stems not merely from technical limitations, but from a deeper mismatch: many leading methods were designed for well-defined, evolutionarily conserved orthosteric sites, whereas allosteric sites are often structurally transient, energetically unstable, and weakly conserved—features that defy conventional detection paradigms.^34^ Recognizing these limitations, a recent illuminating study integrated local binding site information, coevolutionary information, and information on dynamic allostery into a structure-based three-parameter model to identify potentially hidden allosteric sites in ensembles of protein structures with orthosteric ligands.^35^

More recent efforts have sought to hybridize AI with physics-based principles. AlloPred uses normal mode perturbations along with pocket descriptors in a machine learning approach to rank allosteric sites, with a good performance of 28 out of 40 cases.^36^ The Allosite approach, which uses support vector machine (SVM) classifiers of topological and physicochemical characteristics of allosteric and non-allosteric sites, has shown promise in predicting allosteric pockets.^37^ AllositePro, an extension of Allosite, combines pocket features with perturbation analysis to identify allosteric sites and was able to validate a novel allosteric site in cyclin-dependent kinase 2 (CDK2).^38^ DeepAllo combines fine-tuned PLM on AlloSteric Database (ASD)^39^ with FPocket features and shows an increase in prediction performance of allosteric sites over previous studies.^40^

These studies indicated that binding site detection using AI-based approaches must be grounded and interpreted through biophysical principles to establish stronger linkages between prediction performance and understanding of biological mechanisms. To address this, we explored local frustration analysis—a framework rooted in energy landscape theory that quantifies local energetic conflicts arising from competing interactions within a folded protein.^41–46^ While the protein core is typically minimally frustrated—optimized for stability through evolution—functional regions such as binding interfaces, catalytic centers, and allosteric switches are enriched in neutral or highly frustrated residues, which confer conformational plasticity, enable ligand-induced remodeling, and facilitate long-range communication. The frustration survey of the diverse protein folds showed that minimally frustrated residues may be enriched in the protein core, while the exposed flexible binding interfaces are characterized by the highly frustrated patches that can be alleviated upon binding.^44–46^ However, our biophysical studies of different protein families showed that the vast majority of protein residues typically display neutral frustration which implies a moderate degree of stabilization/destabilization for the native interactions as compared to the average energetics induced by conformational or mutational changes in the respective positions.^47–50^. The integration of local energetic frustration with the predictive power of AlphaFold2 (AF2) approaches can be utilized as a physical “guide” to sample the conformational space as local frustration often maps to the high-energy, flexible regions governing allosteric dynamics. By integrating Randomized Sequence Scanning (RASS) adaptation in AF2 predictions of protein kinase conformational ensembles with frustration mapping our recent study revealed why AF2 reliably recovers the minimally frustrated active kinase states but fails to model the highly frustrated inactive conformations that are characteristic of the low-populated, fully inactive ABL kinase form and can define energetically frustrated cracking sites of conformational transitions, presenting difficult targets for AF2 models. The emergence of previously unappreciated connections between local frustration patterns of the functional allosteric states and the performance of the AF2 strongly indicated that the energy landscape frustration can serve as an important interpretability metric of AI-based predictions of protein structures and ensembles.^51^ Integration of energetic frustration analysis and AF2 was explored for predicting protein conformational motions and generation of alternative conformations and transition pathways by progressively enhancing frustration features within the input Multiple Sequence Alignment (MSA) in AF2 architecture^52^. By perturbing the evolutionary signals with these energetic constraints, the model is forced to explore previously hidden regions of the structural landscape in adenylate kinase (AdK) as a model system.^52^ These results suggest that coupling landscape-based frustration analysis with AF2 provides an interpretable and computationally efficient framework for capturing the functional motions essential for allosteric regulation.

In the present study, the pretrained ESM2-650M model^53^ was fine-tuned on the human subset of the LIGYSIS dataset^4,54^ for prediction of protein-ligand binding sites. To adapt the general-purpose encoder for the task of binding site detection, we replaced the original masked language modeling head with a token-level classification head that acts as a projection layer that maps the high-dimensional latent representation of each residue to a scalar probability score for a given protein residue to be part of the binding site. Here, we employed this fine-tuned PLM for prediction of binding sites and structure-based detection P2Rank approach for identification of orthosteric and allosteric binding sites in protein kinases. Subsequently, the results of binding site predictions and performance assessments of PLM and P2Rank were contextualized through large scale local frustration analysis of protein kinase complexes with diverse types of kinase inhibitors. We show that successes and failures of AI-based predictions of the binding sites can be rationalized by energetic landscape analysis. Protein kinases that are regulatory machines characterized by the multiple functional states of the catalytic domain^55–57^ present a class of regulatory switches that operate through dynamic equilibrium between the active and inactive states. In protein kinases, the distinction between orthosteric and allosteric binding sites is fundamental to both their biological regulation and the design of selective drugs. While protein kinases share a highly conserved structural catalytic core and the ATP-binding cleft, the therapeutic promise of allosteric modulators—which bind distal pockets to achieve enhanced selectivity, overcome resistance, or fine-tune activity—has intensified the search for robust, scalable methods to detect these elusive sites. In our study we benchmarked PLM and P2Rank approaches on a meticulously curated dataset of kinase–ligand complexes from Kinase Conformation Resource (KinCoRe), developed by the Dunbrack Lab, which is the comprehensive database for the automated classification of all protein kinase structures.^58–60^ For human protein kinases in complexes with all ligands, the total number of covered kinase genes is 453 of the 518 human kinases. The dataset includes type I orthosteric ATP-competitive inhibitors binding active (DFG-in) state; Type II orthosteric ATP-competitive binding inactive (DFG-out) state; Type III allosteric binding to a pocket proximal (adjacent) to ATP site; and Type IV allosteric binding a pocket distal from ATP site. The dataset reflects DFG-in dominance as about 80% of all kinase structures in the PDB are in the DFG-in state. As a result, human protein kinases in complex with Type I ligands covered 274 kinase genes, while type III and type IV allosteric kinase inhibitors covered only 66 kinase genes.^59^

When fine-tuned for binding site prediction, PLM can achieve remarkable performance on ATP-competitive sites in kinases, where strong evolutionary signals and high residue prevalence enable accurate ranking. However, their performance collapses on allosteric sites, which are sparsely populated (<3% of residues), structurally heterogeneous, and often lack deep coevolutionary signatures. Our results reveal a consistent energetic divide: ATP-competitive inhibitors bind minimally frustrated, evolutionarily constrained regions, while allosteric modulators target mostly neutral-to-high frustrated zones that enable conformational and evolutionary plasticity. Critically, local frustration analysis explains not only where ligands bind, but why certain predictors succeed or fail. The limitations of PLM and P2Rank approaches are not necessarily flaw of the models per se, but a reflection of a fundamental biophysical reality where allosteric sites are not under the same evolutionary pressure as catalytic cores, and their functional relevance is often highly context-dependent and sensitive to specific conformational states rather than sequence alone. Overall, the results of this study suggest that the integration of AI-based tools and PLMs for binding site prediction with the energy landscape analysis can present a biophysics-enabled framework for interpreting predictive divergence, offering a mechanistic path toward building next-generation models that can explicitly utilize energetic constraints to enhance both the accuracy and interpretability of binding site prediction in drug discovery.

## 2. Materials and Methods

### 2.1 Data and ligand-proximal ground truth

The KinCore classification system developed by the Dunbrack Lab provides a rigorous, geometry-based nomenclature for kinase inhibitors.^58–60^ Unlike older systems that relied on qualitative descriptions, KinCore uses specific structural markers—primarily the DFG motif (Asp-Phe-Gly) and the regulatory αC-helix position—to categorize ligands into types I, I.5, II, III, and IV Type I Inhibitors are the most common ATP-competitive ligands. They bind to the kinase in its catalytically active state (DFG-in and αC-in). Type I.5 Inhibitors represent a transition category for inhibitors that bind a DFG-in state but require a shift in the αC-helix to C-out position. They occupy the ATP pocket and extend into the “back pocket” (the space created when the C-helix moves “out”).Type II Inhibitors are ATP-competitive but bind only when the kinase is in its inactive, DFG-out conformation with regulatory αC-helix in “in” or “out” positions. Type III Inhibitors are proximal allosteric inhibitors that are non-competitive with ATP. They bind to a pocket adjacent to the ATP site but do not enter the ATP-binding cleft itself. According to KinCore Criterion the type III ligand must have a minimum distance of > 6.0 Å from the hinge region. It must also make at least three contacts with residues in the “back pocket.” They modulate activity by preventing the closure of the lobes or the positioning of the C-helix, effectively acting as a “wedge.” Type IV allosteric inhibitors are distal allosteric inhibitors that bind to truly distal sites, far from the catalytic machinery. Type IV allosteric modulators can bind to any kinase state, as they do not interact with the active site motifs directly. According to KinCore Criterion, type IV allosteric ligand must be > 6.5 Å away from both the hinge region and the C-helix-Glu(+4) residue (a key conserved salt-bridge residue). The distinction between Type III and Allosteric ligands in this framework primarily comes down to proximity to the active site. For each structure–chain pair, ligand-proximal annotations were derived directly from the mmCIF coordinates. For every ligand specified in the shared spreadsheet, we identified all protein residues whose heavy atoms fell within a distance of 4 Å from any heavy atom of that ligand. When multiple ligands were listed for a structure, we took the union of all contacting residues. Residues not satisfying the 4 Å distance criterion were treated as negatives. If a chain appeared in more than one entry or exhibited inconsistent residue numbering across sources, labels were reconciled by the author-provided numeric residue positions, resolving any conflicts conservatively in favor of positive labels. This procedure defined the ligand-proximal ground truth used throughout all evaluations. The protein kinase structures from the dataset were labeled and discussed in the Results as five distinct categories denoted as *Kinase ALLO*, *Kinase Type I*, *Kinase Type I.5*, *Kinase Type II*, and *Kinase Type III*.

### 2.2 Sequence-Based Finetuning Protein Language Model

The workflow begins with the LIGYSIS dataset, a curated collection of protein–ligand complexes, from which protein sequences are extracted as input (Figure 1). These sequences are processed by a pretrained ESM2-650M transformer architecture^53^ trained using a masked language modeling (MLM) objective with 650 Million parameters, 33 transformer layers and 20 attention heads being trained on UniRef50 database to learn deep evolutionary patterns encoded in protein sequences. Within the ESM-2 suite (which ranges from 8 million to 15 billion parameters), the 650M variant is often considered the “sweet spot” for protein structure modeling because it offers a powerful balance between high-resolution biological insight and manageable computational requirements. Unlike the 3B or 15B versions, the ESM2-650M model can typically be fine-tuned or run for inference on a single consumer-grade or mid-range enterprise GPU (A100 or even a 3090/4090). More particularly, the original classification head for masked language modeling was replaced with a binary classification head, where for each residue, the model predicts a probability of that residue being a part of a binding site or not. The model is repurposed for a downstream task via end-to-end finetuning where the whole protein language model was retrained (or finetuned). Unlike PEFT approach, where one inserts additional weights inside the model while keeping the original weights untouched, in our model we did not insert any additional weights but retrained the original weights. Noticeably, finetuning is generally considered to be a more powerful technique than PEFT, but also more computationally demanding.

**Figure 1.**
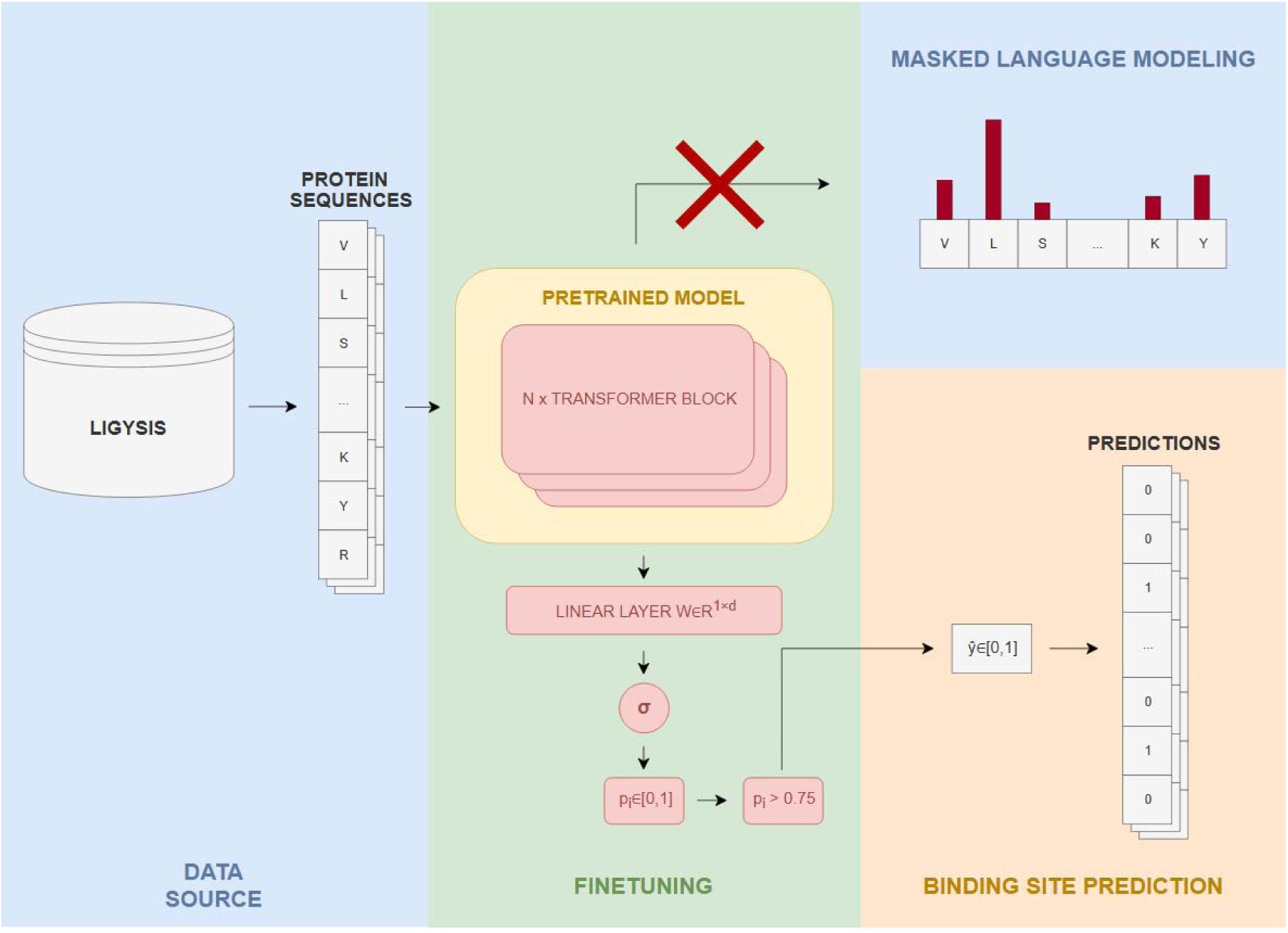
Schematic overview of the fine-tuned protein language model (PLM) pipeline for residue-level binding site prediction. The workflow begins with the LIGYSIS dataset from which protein sequences are extracted as input. These sequences are processed by a pretrained ESM2-650M transformer architecture. The model is repurposed for a downstream task via end-to-end fine-tuning. A new token-level classification head—a single linear layer followed by a sigmoid activation function—is appended to the pretrained encoder. This head maps the high-dimensional latent representation of each residue (of dimension 1280) to a scalar probability score *p_i_* ∊ [0,1], indicating the model’s confidence that residue iparticipates in ligand binding. During finetuning, only the parameters of this new head are initially optimized, after which all weights are jointly refined to ensure compatibility between the learned representations and the task-specific output.

A new token-level classification head—a single linear layer followed by a sigmoid activation function is appended to the pretrained encoder (Figure 1). This head maps the high-dimensional latent representation of each residue (of dimension 1280) to a scalar probability score *p_i_* ∊ [0,1], indicating the model confidence that residue i participates in ligand binding. During fine-tuning, only the parameters of this new head are initially optimized, after which all weights are jointly refined to ensure compatibility between the learned representations and the task-specific output. Final predictions are generated by applying a decision threshold (here, *p_i_* > 0.75) to the predicted probabilities, yielding a binary label (0 = non-binding, 1 = binding) for each residue.

#### 2.2.1 Encoder Representation

Let protein sequence **S** be defined as a sequence of amino acid tokens **S=(x_1_, x_2_, …, x_L_)**, where L is the length of the protein. The sequence is processed by the ESM2-650M encoder, a Transformer-based architecture [<u>10</u>] consisting of 33 layers with 20 attention heads per layer. The encoder maps the discrete input tokens to a sequence of continuous embedding vectors:

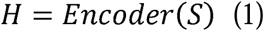

where *H* ∊ *R^L×d^* represents the final hidden state, and **d** is the dimension (d=1280). Each vector *h_i_* ∊ *R^d^* in **H** contains the contextualized representation of the residue at position i, aggregating information from the entire sequence via the self-attention mechanism.

#### 2.2.2 Fine-tuning Binding Site Projection Head

To adapt the general-purpose encoder for the task of binding site detection, we replaced the original masked language modeling head with a token-level binary classification head. This head acts as a projection layer that maps the high-dimensional latent representation of each residue to a scalar probability score. Specifically, the high-dimensional vector h_i_, which encodes the residue’s local chemical environment and global evolutionary context, is transformed into a single scalar logit. This logit is then passed through a Sigmoid activation function to produce a probability score *p_i_* ∊ [0,1], representing the model’s confidence that the specific residue participates in ligand binding.

For each residue i, the probability p_i_ of the residue being part of a binding site is computed as:

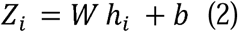

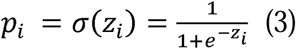

Here, *W* ∊ *R*^1×*d*^ is the learnable weight matrix of the binary classification head, *b* ∊ *R*^1^is the bias term, σ denotes the Sigmoid activation function and p_i_∈[0,1].

#### 2.2.3 Optimization and training

The model was trained to minimize the weighted Binary Cross-Entropy (BCE) loss:

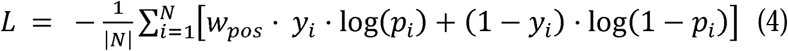

where *y_i_* ∊ {0,1} is the ground truth label for residue i (1 for binding site, 0 otherwise), and w_pos_ is a weighting factor introduced to counteract class imbalance. The model was optimized using the AdamW optimizer^61^ with a learning rate of 1e−4 and L2 regularization.

The fine-tuning ran for 3 epochs. Similar to other studies^29^, to stabilize the fine-tuning, the newly initialized binary classification head was trained during the first two epochs, while the rest of the model was frozen. After this warm-up phase, all weights were unfrozen and finetuned jointly. Due to memory constraints, the training was performed with a batch size of 1. The decision threshold of 0.75 was used as a threshold to distinguish between positive and negative classes. This threshold was based on maximizing the Matthews Correlation Coefficient (MCC). MCC is a powerful metric for evaluating binary classification models, known as a balanced measure, even with imbalanced datasets, ranging from –1 (perfect misclassification) to +1 (perfect classification), with 0 being random guessing, calculated using true positives, true negatives, and false negatives from a confusion matrix.^62^ It provides a comprehensive score by considering all four confusion matrix categories. The model was fine-tuned on the human subset of the LIGYSIS dataset.^4,54^ To set the parameters (learning rate, dropout, number of epochs), a validation subset was selected randomly from the LIGYSIS dataset. MCC was utilized for determining optimal parameters. After setting the parameters, the whole LIGYSIS dataset (including the validation subset) was utilized to fine-tune the final model.

#### 2.2.4 Sequence similarity control

Because the PLM was pre-trained and fine-tuned on a large collection of protein sequences that may have included many kinases, we controlled for possible train–test leakage by removing all evaluation chains whose sequences overlapped with those used for PLM training. For each chain in the kinase dataset, we used its UniProt identifier as provided in the curated metadata. We then compared these identifiers against the list of UniProt entries that had been used during PLM fine-tuning. Any chain whose UniProt accession appeared in the PLM training list was considered potentially exposed to the model and was excluded from the evaluation. This filtering strategy functioned as a conservative safeguard: chains were removed whenever they shared the same UniProt accession with the training set, regardless of isoforms or engineered sequence variants.

All results reported in this study were therefore computed on the remaining set of non-overlapping chains, so that the evaluation reflected genuine generalization rather than memorization of training sequences. Retention varied across kinase families: the allosteric set retained the largest fraction of chains (54.2 %), while the Type III family retained the fewest (30.3 %). Type I, Type I.5, and Type II showed intermediate retention values of approximately 35–38 %. Detailed per-family counts and percentages are summarized in Table 1. The resulting checkpoint outputs, for each residue, a probability *p_i_* ∊ [0,1] that the residue belongs to a ligand-proximal site. In the final PLM model, there is only the single head for the binding site prediction.

**Table 1.** Retention after removal of chains whose UniProt accessions overlapped with those used during PLM fine-tuning. Each row corresponds to a PDB chain entry in the kinase dataset.

| Dataset | Total chains | Overlapping with PLM | Retained chains | Retention (%) |
| --- | --- | --- | --- | --- |
| Kinase ALLO | 675 | 309 | 366 | 54.2 |
| Kinase Type I | 7604 | 4713 | 2891 | 38.0 |
| Kinase Type I.5 | 775 | 507 | 268 | 34.6 |
| Kinase Type II | 553 | 361 | 192 | 34.7 |
| Kinase Type III | 294 | 205 | 89 | 30.3 |

In the *PLM fixed* setting, the probabilities were binarized using a single global threshold *t* = 0.75. In the *PLM bestMCC* setting, we selected one global threshold *t*\* by sweeping *t* ∊ [0,1] in increments of 0.05 and choosing the value that maximized the micro-averaged MCC on the evaluation set. Raw probabilities were retained for threshold-free analyses using AUROC and AUPR. Inference was performed using custom Python scripts that wrap the fine-tuned ESM2 checkpoint and implement residue extraction, tokenization, batching, and model evaluation.

### 2.3 Structure-Based P2Rank Approach

P2Rank (version 2.5) was used as a structure-based baseline.^10,11^ For each mmCIF structure P2Rank predicted one or more putative ligand binding pockets with residue-level scores. To obtain per-residue predictions compatible with the PLM and with the ligand-proximal ground truth, we used the “ANY-pocket” aggregation: each residue was assigned the maximum probability over all predicted pockets that contained that residue. If a residue was absent from all pockets, we assigned a score of 0. We then evaluated P2Rank at its default per-residue decision threshold under this aggregation, which we refer to as *P2 ANY fixed*. Where informative, we also considered threshold sweeps analogous to the PLM setting, but the main comparisons are reported for the default operating point. P2Rank is a stand-alone command-line program for fast and accurate prediction of ligand-binding sites from protein structures.^4,10,11^

### 2.4 Evaluation protocol

All computations respected the structure–chain key (*PDBID, chain*). Predictions from the PLM and P2Rank were aligned to the ground truth by numeric residue indices using author numbering extracted from the mmCIF files. Only residues belonging to the evaluated chain and present in all three sources (ligand-proximal ground truth, P2Rank outputs, PLM predictions) were included in the analysis. Dedicated validation scripts confirmed that, after dataset filtering, residue indices and residue counts matched exactly across all sources. We report on both micro-and macro-averaged performance measures. In the *micro* setting, residues from all chains were concatenated before computing a metric, emphasizing overall operating behavior and the effects of class imbalance at the dataset level. In the *macro* setting, metrics were computed independently for each chain and summarized by the mean, with variability reported as half of the interquartile range (IQR/2), reflecting per-structure robustness.

Accuracy and MCC metrics followed standard definitions. The F_1_ score was the weighted variant, averaging per-class F_1_ values according to class support. From raw PLM probabilities we also computed AUROC, which measures the probability that a randomly chosen positive residue ranks above a randomly chosen negative and is comparatively insensitive to prevalence, and AUPR, which summarizes the precision–recall trade-off and is more informative under the strong class imbalance typical of residue-level binding-site prediction.

### 2.5 Local Frustration Analysis

We explored local frustration analysis which is a framework rooted in energy landscape theory that quantifies local energetic conflicts arising from competing interactions within a folded protein.^40–42^ Configurational frustration evaluates the sensitivity of the interaction to local structural perturbations by randomizing both residue identities and interatomic distances within the native contact geometry. Mutational frustration assesses the energetic optimality of a native amino acid pair (i, j) by comparing its interaction energy to that of all viable alternative amino acid pairs at the same structural positions, while keeping the backbone geometry fixed. The local energetic frustration index is computed as a Z-score using the contribution of a residue to the energy in a given conformation as compared to the energies that would be found by mutating residues in the same native location (mutational frustration) or by creating by changing local conformational environment for the interacting pair (conformational frustration). A residue-based frustration index measures the energetic stability of a particular native contact as compared to a set of all contacts sampled by generating an ensemble of 1,000 distributed decoys and recomputing the energy change. Following the established validated criteria for differentiating frustration categories across a wide range of biomolecular systems^40–42^, the contacts were classified as minimally frustrated if Z > 0.78 (native interaction is significantly more favorable than alternatives), highly frustrated if Z < −1.0 (native interaction is significantly less favorable), and neutrally frustrated otherwise (native interaction is neither strongly favored nor disfavored). To assign a residue-level frustration profile, we computed the local density of frustrated contacts within a 5 Å radius of each residue. A residue was classified based on the dominant frustration type among its interacting partners that appeared in >50% of the structural ensemble (i.e., across the most populated conformational clusters).

## 3. Results and Discussion

### 3.1 PLM and P2Rank Benchmarking of Binding Site Detection in Kinase Families Reveals Fundamental Divergence Between Predictions of Orthosteric and Allosteric Sites

To evaluate the capabilities of modern AI-driven binding site predictors in a therapeutically critical protein family, we benchmarked PLM against the structure-based method P2Rank across a biophysically curated dataset of human kinase–ligand complexes. Our analysis spans five functional classes—Type I, Type I.5, and Type II (orthosteric ATP-competitive inhibitors) and Type III and ALLO (allosteric modulators). We employed multiple complementary performance metrics to dissect both global ranking behavior and operational precision.

We first assessed PLM performance using threshold-free metrics: the area under the receiver operating characteristic curve (AUROC) and the area under the precision–recall curve (AUPR). AUROC reflects a model’s ability to rank true positives above negatives, regardless of class balance, while AUPR is extremely sensitive to sparsity and reflects operational utility in real-world screening. As shown in Figure 2A, the PLM achieves excellent AUROC across all orthosteric families: 0.965 (Type I), 0.974 (Type I.5), and 0.952 (Type II). This indicates that, even without tuning a decision threshold, the model consistently assigns higher binding probabilities to residues within the ATP-binding cleft than to non-binding surface residues. In contrast, allosteric sites (Type III and ALLO) show markedly lower AUROC (0.891 and 0.705, respectively) suggesting that while the model retains some ranking signal, the separation between true and false positives is less distinct. The analysis of the AUPR metric (Figure 2B) reveals a more detailed picture, collapsing dramatically for allosteric sites (0.288 for Type III and a near-baseline 0.061 for ALLO) as compared to 0.640–0.755 for orthosteric kinase complexes. Strikingly, the allosteric set displayed both lower AUROC (0.70) and extremely low AUPR (0.06), consistent with sparse and heterogeneous allosteric sites that are difficult for the model to detect. The Type III family occupied an intermediate regime: AUROC remained high (0.89), suggesting that positives still tended to rank above negatives overall, but AUPR dropped to 0.29, reflecting reduced precision among the highest-ranked residues under strong class imbalance. This stark contrast highlights a critical insight indicating that ranking ability does not guarantee actionable precision. Even if allosteric residues are ranked above random negatives (moderate AUROC), they are drowned in a sea of false positives. This performance gap is strongly correlated with the extreme sparsity (<3% of residues) of allosteric sites and the lack of conserved sequence motifs.

**Figure 2.**
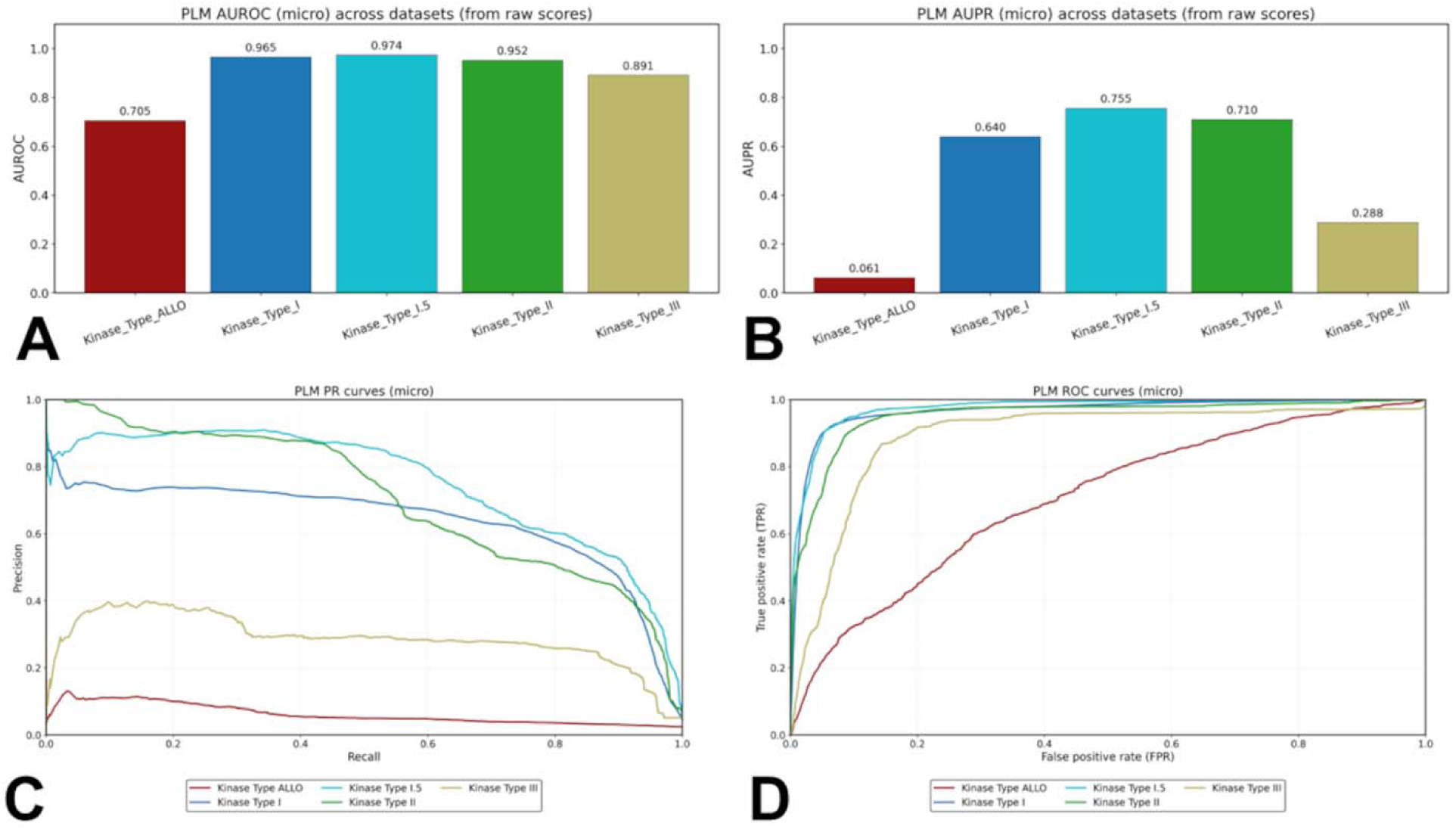
Threshold-free ranking performance of the fine-tuned PLM across kinase inhibitor classes. (A) Micro-averaged AUROC of PLM residue-level scores across kinase families on the evaluation set. Bars report AUROC computed from raw probabilities, with colors corresponding to Kinase Type ALLO (red), Type I (blue), Type I.5 (cyan), Type II (green), and Type III (khaki). (B) Micro-averaged AUPR of PLM residue-level scores across kinase families on the same evaluation set. AUPR is computed from raw probabilities under the same color scheme and highlights the impact of class imbalance: Type I/Type I.5/Type II show consistently high precision–recall performance, whereas allosteric and Type III sites are markedly harder. (C) Precision–recall curves (micro-averaged) for raw PLM probabilities across kinase families. (D) ROC curves (micro-averaged) for raw PLM probabilities across kinase families. Orthosteric families (Type I, I.5, II) exhibit high performance in both metrics, whereas allosteric families (Type III, ALLO) show a pronounced drop in AUPR despite moderate AUROC highlighting the challenge of precision under sparsity.

However, we must distinguish between the observed correlation and potential unique causation. While the lack of conserved motifs is an exacerbating factor, the reduced precision in predicting allosteric binding sites may also stem from structural heterogeneity and context-dependent plasticity inherent to allosteric pockets, which may limit sequence-based predictability, regardless of motif conservation. Notably, PLM embeddings retain sensitivity for detecting remote homology in other contexts, suggesting that the challenge here stems not from an inherent model limitation, but from fundamental differences in how evolutionary signals are encoded at allosteric versus orthosteric sites. This distinction implies that the sequence heterogeneity of allosteric site residues may be a symptom of functional plasticity and conformational ambiguity rather than its root cause—a hypothesis we evaluate directly through frustration analysis in subsequent sections.

Across kinase families, the ranking performance of the PLM showed a clear systematic pattern (Table 2, Figure 2). For Type I, Type I.5, and Type II kinases, the model achieved very high AUROC values (0.95–0.97) together with relatively high AUPR (0.64–0.76), indicating that residues from the ligand-proximal class were consistently ranked above negatives and that many of the top-scoring residues corresponded to true binding-site positions. The results suggest that the divergence between AUROC and AUPR for type III and type IV allosteric sites may not only point to some limitations of the PLM but more significantly present a symptom of the inherent semantic ambiguity of these regions in sequence space.

**Table 2:**
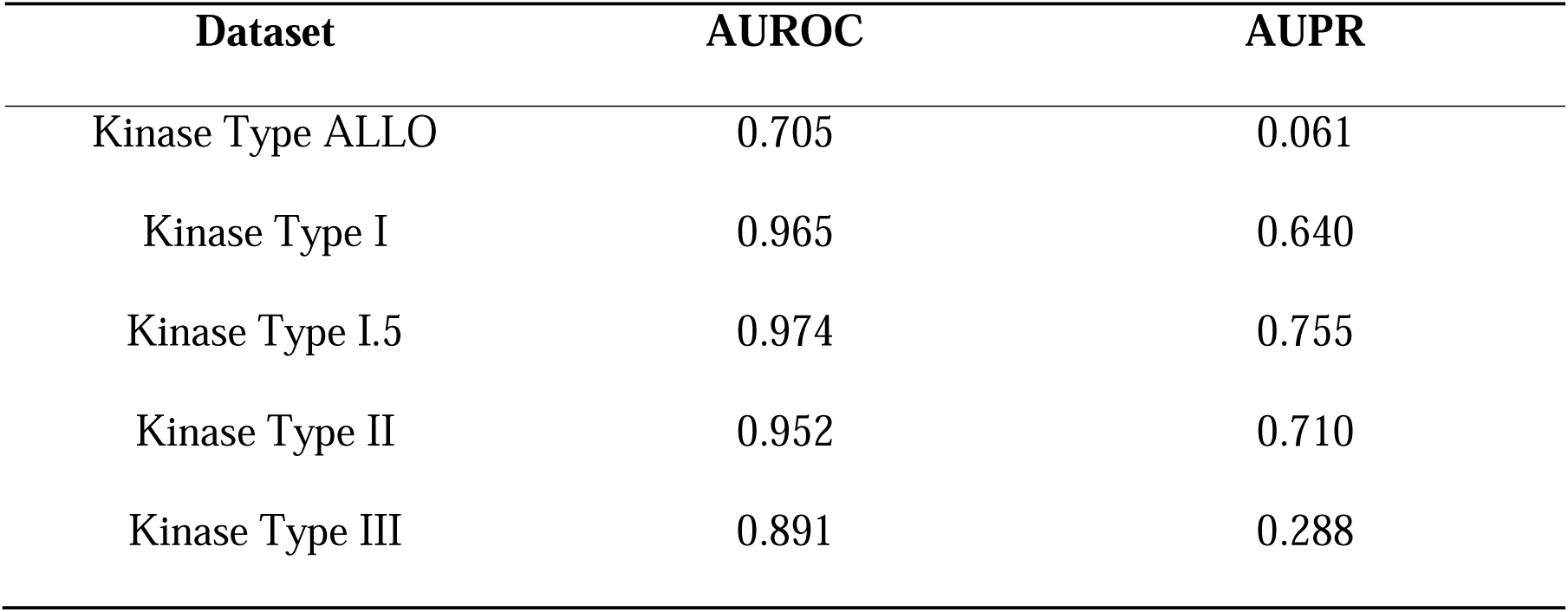
Micro-averaged AUROC and AUPR computed from raw PLM probabilities on the filtered kinase evaluation sets.

To further dissect this behavior, we examined full precision–recall (PR) and ROC curves (Figure 2C,D). The PR curves for Type I–II inhibitors showed that precision remains high (>0.7) even as recall increases, reflecting a dense, conserved signal in the ATP cleft (Figure 2C). In contrast, for ALLO and Type III, precision plummets as soon as recall exceeds 0.1, indicating that the top-ranked residues are dominated by false positives (Figure 2C). The ROC curves (Figure 2D) remain high across all families, confirming that the PLM maintains strong global discriminative power.

This behavior is characteristic of extreme class imbalance combined with low intrinsic signal quality. Notably, Type III complexes (0.288 AUPR) perform better than ALLO class (0.061), consistent with its slightly higher prevalence (4.77% vs. 2.96%) and proximity to the ATP site.

To contextualize the difference between AUROC and AUPR Table 3 reports the number of positive and negative residues per family on the filtered evaluation set (positives are residues with any heavy atom within 4 Å of an annotated ligand; all others are negatives).

**Table 3:** Class balance on the filtered evaluation set.

| Dataset | Chains | Positives | Negatives | Prevalence (%) |
| --- | --- | --- | --- | --- |
| Kinase Type ALLO | 849 | 8140 | 267118 | 2.96 |
| Kinase Type I | 3349 | 48241 | 1002961 | 4.59 |
| Kinase Type I.5 | 277 | 5136 | 79930 | 6.04 |
| Kinase Type II | 209 | 4233 | 55646 | 7.07 |
| Kinase Type III | 117 | 1644 | 32833 | 4.77 |

As shown in Table 3, the allosteric category contains by far the lowest fraction of positive residues (2.96%). With 8,140 positives and 267,118 negatives, the extreme class imbalance explains why AUROC remains moderately high (0.705) while AUPR collapses to 0.061: even a well-ranked prediction list yields low precision when positives are this sparse. Type III family shows a similar but less extreme pattern, with prevalence of 4.77%, which contributes to its intermediate AUPR of 0.288 despite a high AUROC of 0.891. In contrast, Type I, Type I.5 and Type II families show substantially higher prevalence (4.59–7.07%), directly supporting their much higher AUPR (0.64–0.76). The ordering of difficulty across kinase families aligned closely with both biochemical expectations and the underlying class balance. Type I and Type I.5 inhibitors bind in the ATP cleft, which contains strongly conserved sequence motifs. Their high AUROC values (0.96–0.97) and elevated AUPR (0.64–0.76) indicate that the PLM effectively captures these conserved binding determinants using sequence-derived representations alone. Type II inhibitors extend from the ATP site into the back cleft and depend on the DFG-out conformation. Their AUPR (0.71) is slightly lower than for Type I.5 but remains comparable to or higher than Type I, consistent with a moderate increase in conformational diversity while retaining substantial sequence signal. In all three ATP-site families, prevalence is relatively high (4.6–7.1%), supporting better precision–recall behavior. Type III inhibitors bind distal, partially cryptic pockets, and the Allosteric dataset consists entirely of non-ATP allosteric sites. These pockets are more heterogeneous, often located in flexible regions, and exhibit much lower prevalence (4.8% for Type III and only 2.96% for allosteric). As a result, AUPR and MCC decrease sharply in these families even though AUROC remains moderate to high (0.89 for Type III and 0.71 for Allosteric). This reflects the well-known behavior of precision–recall metrics under severe class imbalance: even when positives are ranked above negatives on average, precision drops rapidly when positives are sparse. These trends are corroborated by the ROC and PR curve overlays (Figure 2C,D). ROC curves remain high for all families, demonstrating reliable global ranking of positives versus negatives. In contrast, PR curves show a pronounced decline in precision for Type III and especially allosteric binding sites as recall increases, capturing both their greater structural heterogeneity and the extreme imbalance of binding versus non-binding residues.

### 3.2 PLM Outperforms P2Rank on Orthosteric Sites Across Micro-Averaged and Macro Metrics

Next, we compared the PLM against P2Rank, a leading structure-based method that predicts “ligandability” from surface geometry and chemical features. To ensure fair comparison, P2Rank predictions were aggregated using the “any-pocket” strategy, assigning each residue the maximum score across all predicted pockets. At the micro level (aggregating all residues across all structures), the PLM outperforms P2Rank on orthosteric families across all metrics (Figure 3). For MCC metric a balanced measure of classification quality, the PLM achieves 0.58–0.69, compared to 0.42–0.54 for P2Rank (Figure 3A).

**Figure 3.**
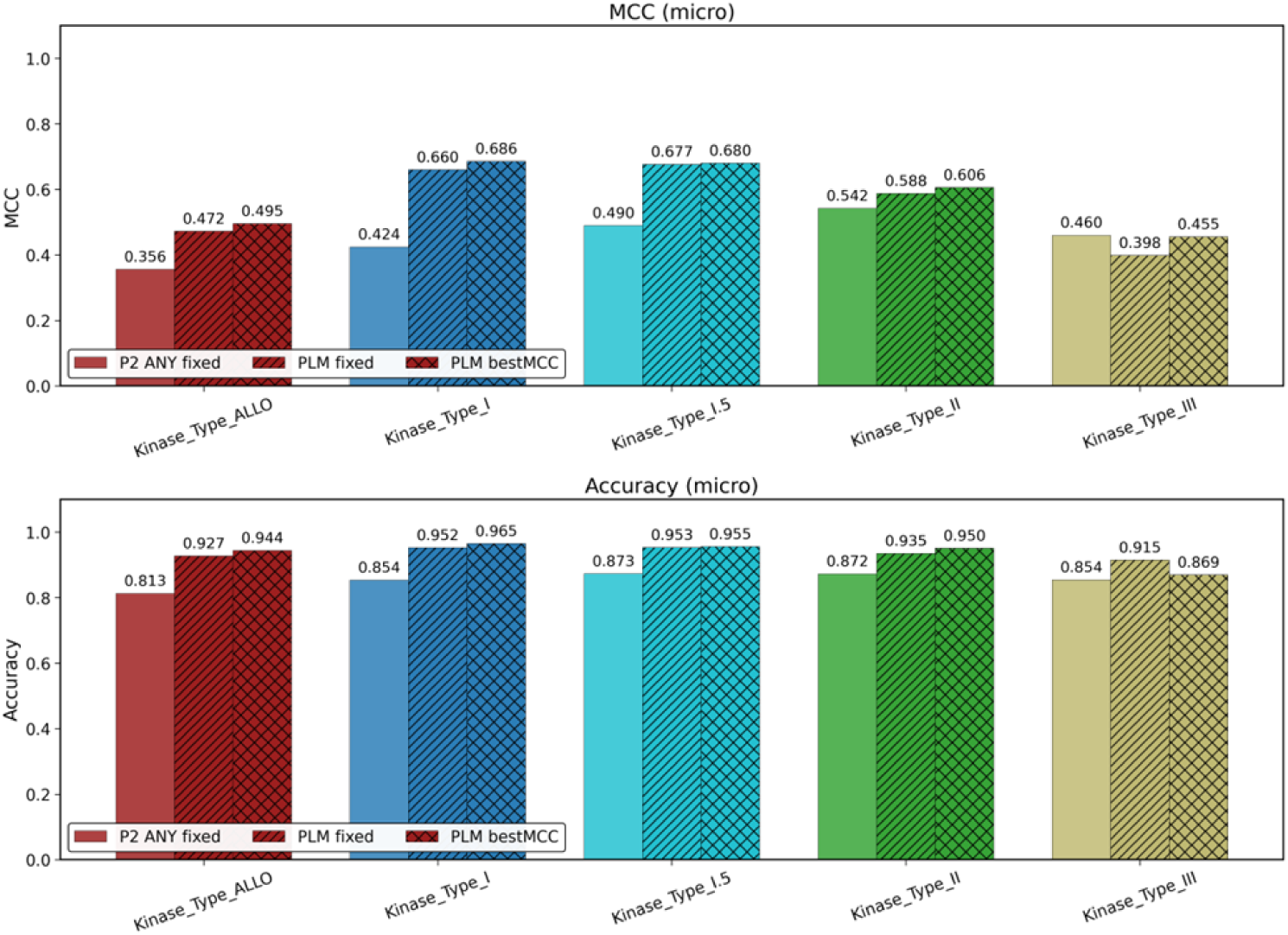
The distribution of micro-averaged metrics across diverse types of protein kinase complexes. (A) MCC metric and (B) accuracy across kinase families. The PLM (blue) consistently exceeds P2Rank (orange) for Type I–II inhibitors, with smaller gains for Type III and ALLO—reflecting the intrinsic difficulty of predicting evolutionarily labile sites.

Similarly, accuracy rises from 0.85–0.87 (P2Rank) to 0.94–0.96 (PLM) (Figure 3B). These gains reflect the PLM’s ability to integrate evolutionary context beyond static structural features. For allosteric families, the gap narrows: PLM and P2Rank achieve comparable MCC (0.44–0.47 vs. 0.35–0.47), underscoring that both approaches struggle when functional sites lack strong signals—whether evolutionary (PLM) or geometric (P2Rank).

To ensure that our results were not driven by a few dominant or well-behaved structures, we computed macro-averaged metrics, which average performance per chain before aggregating (Figure 4).

**Figure 4.**
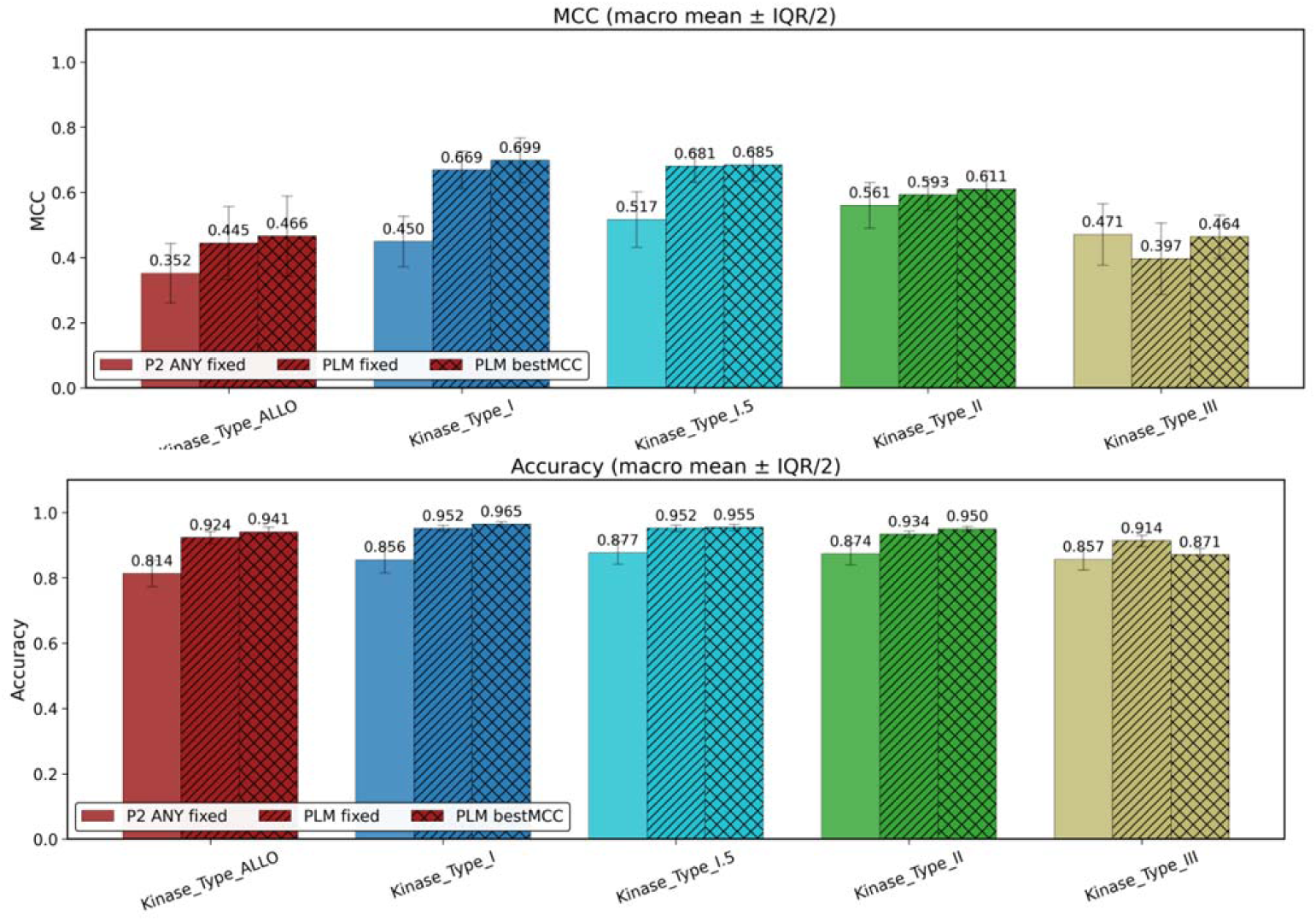
The distribution of macro-averaged metrics across diverse types of protein kinase complexes. (A) MCC and (B) accuracy, summarized as mean ± half-IQR across individual kinase structures.

This approach reveals per-structure robustness and guards against dataset skew. The pattern mirrors the micro analysis: for Type I and I.5 kinases, macro MCC increases from ∼0.48 (P2Rank) to ∼0.69 (PLM), with similarly strong gains in accuracy (0.87 → 0.96). For the allosteric and Type III families the separation was more modest, with PLM achieving 0.44–0.47 compared with 0.35–0.47 for P2Rank, reflecting their higher structural heterogeneity and lower prevalence of contacting residues. Together with reduced macro variability, this indicates that the PLM improves performance across most structures, not only the best cases. Crucially, the interquartile range (IQR) is narrower for the PLM, indicating lower variability across diverse kinase scaffolds. This demonstrates that PLM improvements are systematic, not confined to a subset of well-conserved kinases. For allosteric kinase complexes, both methods show higher variability and reduced performance, reflecting structural heterogeneity of allosteric pockets. Finally, we evaluated F1 score, the harmonic mean of precision and recall (Figure 5).

**Figure 5.**
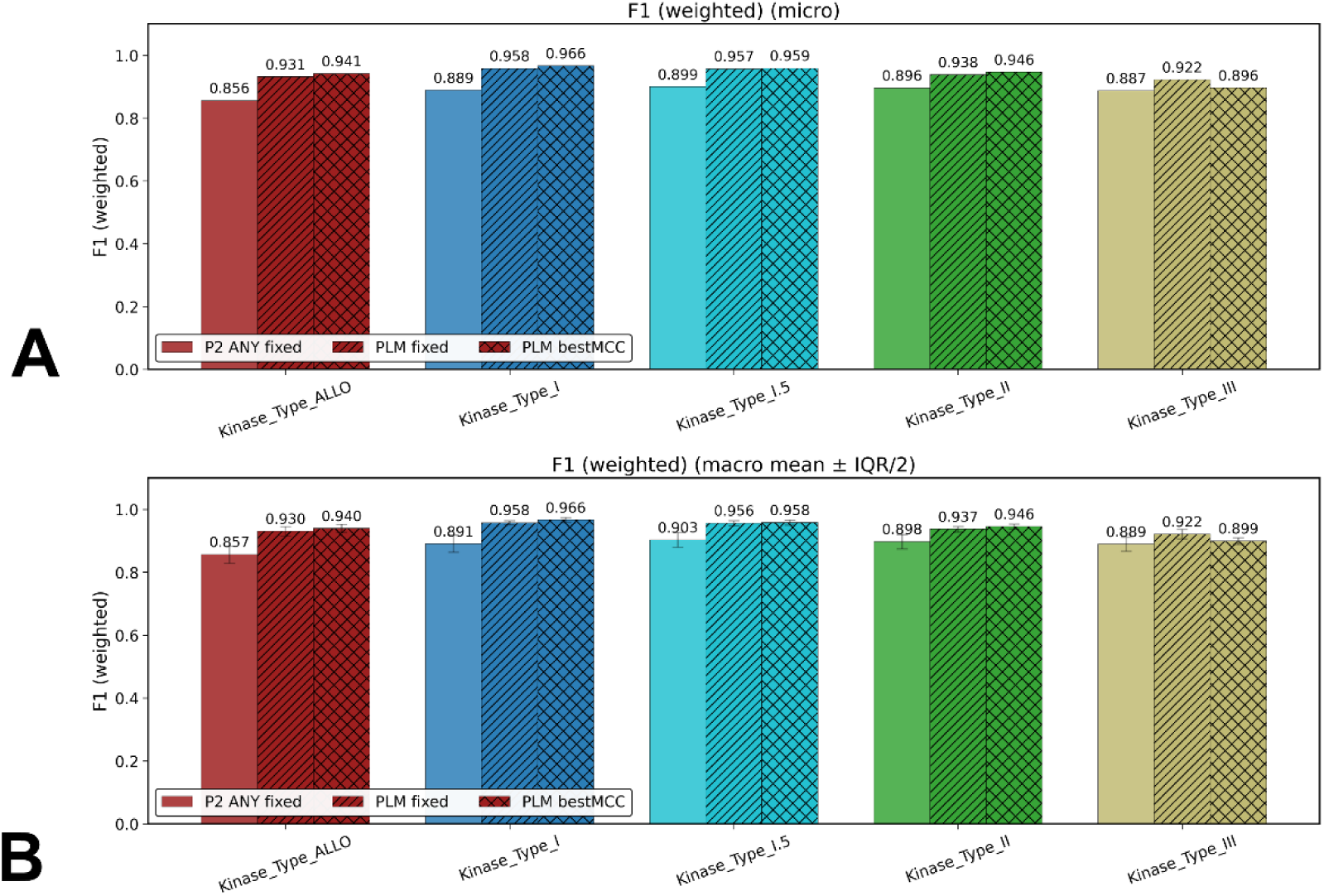
(A) Micro F_1_ (weighted) across kinase families. (B) Macro F_1_ (weighted) across kinase families. Error bars show IQR/2.

Micro-averaged and macro-averaged F1 scores reinforce the central finding that PLM excels where evolutionary signals are strong (orthosteric sites) but plateau where they are weaker (allosteric sites). Micro-averaged F1 shows the same trend: 0.96–0.97 for orthosteric PLM predictions, versus 0.89–0.90 for P2Rank, with smaller but consistent gains for allosteric binding sites (0.78–0.82 vs. 0.72–0.77). where sparse positives limit the achievable F1 regardless of method. Macro-averaged F1 values confirmed that the improvement is systematic and not driven by a small number of structures (Figure 5B). Notably, the F1 gap is smallest for ALLO category, where both precision and recall are inherently limited by sparsity.

In summary, the results revealed that orthosteric sites (Type I/II) are predicted with high accuracy, precision, and robustness by the PLM, reflecting their strong evolutionary conservation and high prevalence (4.6–7.1% of residues). At the same time, allosteric sites (Type III/ALLO) can suffer from catastrophic precision collapse (AUPR < 0.3), not because the model cannot rank them (AUROC remains moderate), but because they lack discriminative sequence signatures in a background of neutral surface residues. Our findings suggest that class imbalance contributes but does not uniquely define the outcome of predictions. Indeed, type III sites (4.77% prevalence) still show poor AUPR (0.29), while Type I.5 (6.04%) achieves high AUPR (0.76) indicating that signal quality matters more than prevalence alone. Finally, PLM consistently outperforms P2Rank on orthosteric sites, but both methods can struggle on allosteric pockets, suggesting that neither sequence nor static structure alone is sufficient for robust and consistent allosteric site detection. We argue that an important and often overlooked issue is semantic ambiguity as allosteric sites lack conserved sequence motifs, rendering them indistinguishable from neutral surface residues in sequence space. Together, these findings establish a performance baseline for binding site prediction in kinases and expose limitations of PLM and P2Rank approaches in detecting evolutionary plastic, context-dependent pockets. The findings also suggest that a performance gap in prediction of orthosteric vs allosteric binding sites that cannot be solely explained by data sparsity alone but instead points to fundamental differences in the evolutionary and energetic signatures of the binding sites.

### 3.3. Local Frustration Analysis Reveals Landscape-Encoded Determinants of Predictive Divergence in Kinase Binding Sites

When fine-tuned for binding site prediction, PLMs achieve remarkable accuracy on conserved orthosteric sites, such as the ATP cleft in kinases, where strong evolutionary signals exist. However, their performance degrades dramatically on allosteric sites, which are often non-conserved, structurally heterogeneous, and sparsely represented in sequence space. Benchmarking studies reveal a consistent pattern: PLMs yield high AUROC and AUPR for Type I/II kinase inhibitors (AUPR > 0.7) but collapse on allosteric datasets (AUPR < 0.1), not due to poor ranking, but because true positives constitute <3% of residues—rendering precision impossible to achieve without additional biophysical context.

We conjectured that this disconnect between prediction performance and biological function is not entirely a result data sparsity but is a more fundamental consequence of the intrinsic biophysical design of allosteric sites. To address this and mechanistically explain the divergent performance of PLMs on orthosteric versus allosteric kinase binding sites, we explored local frustration analysis—a framework rooted in energy landscape theory. By performing a systematic local frustration analysis across a curated set of human kinase–ligand complexes, stratified by ligand type, we characterized key landscape-based signatures of orthostatic and allosteric kinase binding sites. Frustration analysis partitions residue–residue interactions into three energetic classes—minimally, neutrally, and highly frustrated—based on how the native interaction energy compares to an ensemble of decoys generated either by randomizing residue identities (mutational frustration) or by perturbing both identities and local geometry (conformational frustration). Our analysis quantifies both conformational frustration (sensitivity to structural perturbations) and mutational frustration (sensitivity to amino acid substitutions) using the Frustratometer approach^40–42^ which computes residue-level energetic Z-scores relative to decoy ensembles generated by randomizing either residue identities (mutational) or local geometry (conformational).

While the protein core is typically minimally frustrated—optimized for stability through evolution—functional regions such as binding interfaces and allosteric switches can be neutral or highly frustrated residues, which confer conformational plasticity, enable ligand-induced remodeling. We quantified conformational and mutational frustration of protein residues in the protein kinase structures separating kinase complexes into two major categories: type I orthosteric ATP-competitive inhibitors binding active (DFG-in) state and Type II orthosteric ATP-competitive binding inactive (DFG-out) state form first group, while Type III allosteric binding to a pocket proximal to ATP site and Type IV allosteric binding a pocket distal from ATP site formed the second category. Using recent comprehensive survey of allosteric kinase ligands, we similarly mapped the 12 previously characterized allosteric binding sites.^63,64^ (Figure 6).

**Figure 6.**
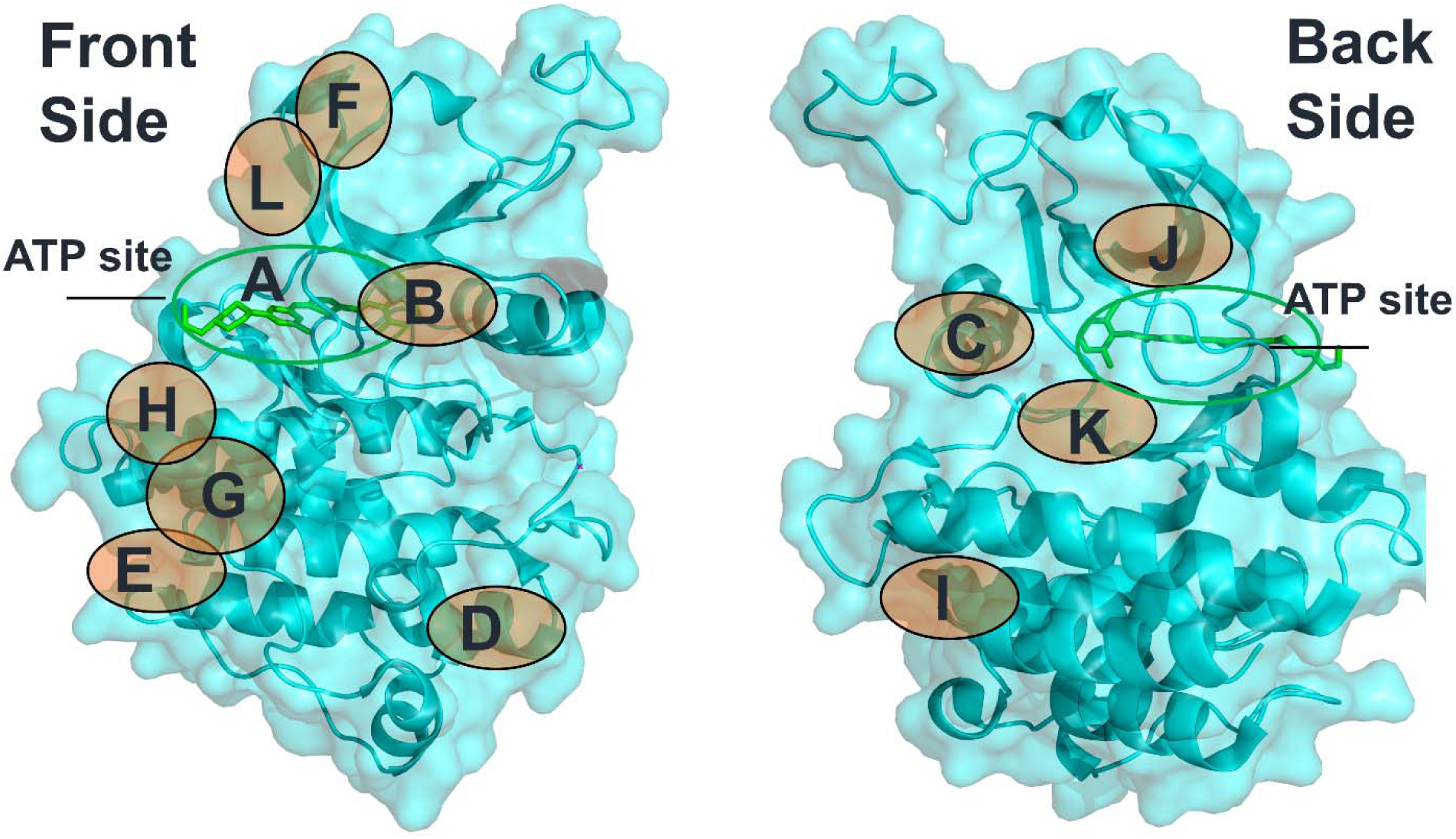
Structural mapping of binding sites for allosteric protein kinase ligands. The distribution of 262 allosteric kinase ligands (including multi-site ligands) over the 12 binding sites mapped on a representative catalytic domain of ABL kinase complex with Dasatinib (pdb id 2GQG). The mapping mirrors a similar analysis based on survey of 262 allosteric kinase ligands and 12 validated allosteric binding sites in protein kinases.^63,64^

We started by computing the cumulative distribution of the conformational and mutational frustration residue indexes across protein kinase complexes of these two major categories. For each residue, we calculated its local frustration level using the Frustratometer framework^41,42^ which evaluates the energetic favorability of native residue–residue contacts relative to decoy ensembles generated by either (i) randomizing interacting amino acid identities (mutational frustration) or (ii) perturbing both identities and inter-residue distances (conformational frustration). Residues were then classified as minimally frustrated (Z > 0.78), neutral (−1 ≤ Z ≤ 0.78), or highly frustrated (Z < −1), following established Frustratometer V2 criteria. The residue distributions showed the overwhelming preferences for conformational neutral and minimal frustration interactions in the protein kinase catalytic domain complexes (Figure 7A,B). Strikingly, the global conformational frustration profiles of orthosteric kinase complexes and allosteric kinase complexes were nearly indistinguishable in their overall architecture (Figure 7A,B). A similar distribution pattern was seen for mutational frustration indexes (Figure 7C,D) showing that a significant number of the kinase domain residues are characterized by relatively high neutral frustration values, while minimal frustration profile spans a wide range of values between 0.2-0.6.

**Figure 7.**
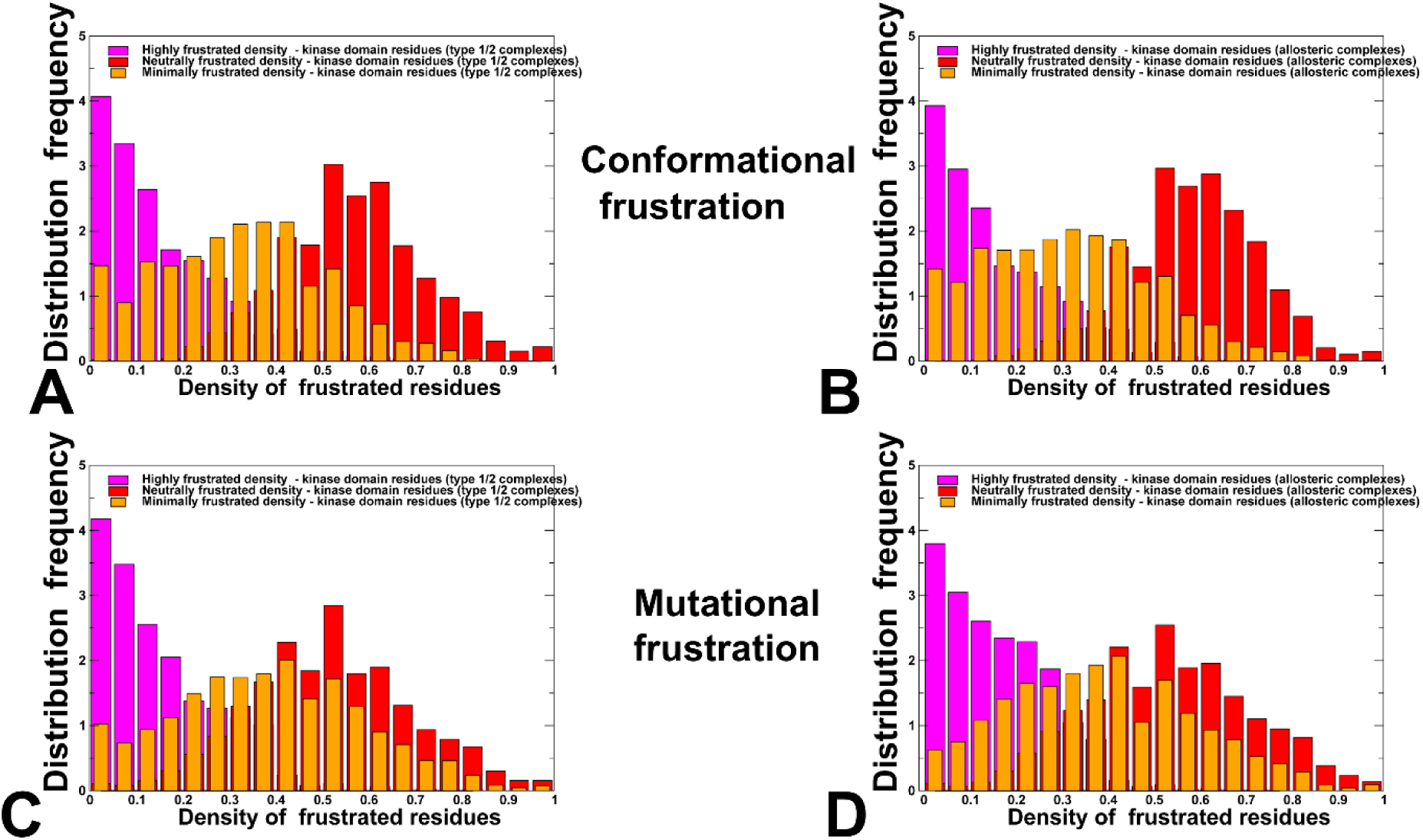
The distributions of conformational and mutational frustration in the protein kinase complexes with type 1/II kinase inhibitors binding to the orthosteric ATP binding site. (A,C) and protein kinase complexes with type III/IV kinase inhibitors binding to diverse allosteric sites. The relative global densities of conformational frustration for the orthosteric kinase complexes (A) allosteric kinase complexes (B). The relative global densities of mutational frustration for the orthosteric kinase complexes (A) and allosteric kinase complexes (B). The highly frustrated density of kinase domain residues is shown in magenta-colored filled bars, neutrally frustrated density is in red-colored bars and minimally frustrated density is in orange-colored bars.

Critically, the similarity in both conformational and mutational frustration distributions across Type I/II orthosteric and Type III/IV allosteric kinase complexes indicates that the global energetic architecture of the kinase catalytic domain is largely conserved, irrespective of ligand binding mode. These results suggest that kinase regulation—whether orthosteric or allosteric—operates within a shared, evolutionarily encoded framework of conformational plasticity, where neutral frustration predominates and enables low-energy transitions between active and inactive states. This topological and dynamic neutrality also extends to sequence variation: many residues tolerate benign mutational changes without compromising fold integrity. However, this mutational tolerance can be more region-dependent; while much of the kinase surface exhibits neutral mutational frustration, the orthosteric ATP-binding site displays pronounced minimal mutational frustration that reflects its essential catalytic role and limited tolerance for substitutions. This regional constraint underlies the higher sequence conservation of the active site and explains the divergent performance of sequence-based predictors on orthosteric versus allosteric pockets, as further dissected in the mutational frustration analysis below.

Highly frustrated residues are sparse, comprising <15% of the domain and exhibiting low relative densities (∼0.0–0.15). They are localized in the C-lobe including portion of the P+1 loop and the αG-helix which serves as a scaffold for activation loop (A-loop) docking in inactive conformations (Figure 8A). In the fully inactive ABL state (pdb id 3K5V) the A-loop folds onto the αG-helix and together with P+1 loop forms a contiguous high frustrated conformational cluster. This region is neutrally frustrated mutationally—evolutionarily permissive—but functionally essential for autoinhibition. The structural map of highly frustrated sites (Figure 8A) reflects concentration of corresponding hotspots in this region of C-lobe. The high frustration residues also occupy a localized region in the N-lobe near regulatory C-helix that orchestrates conformational switches between the inactive and active states (Figure 8A).

**Figure 8.**
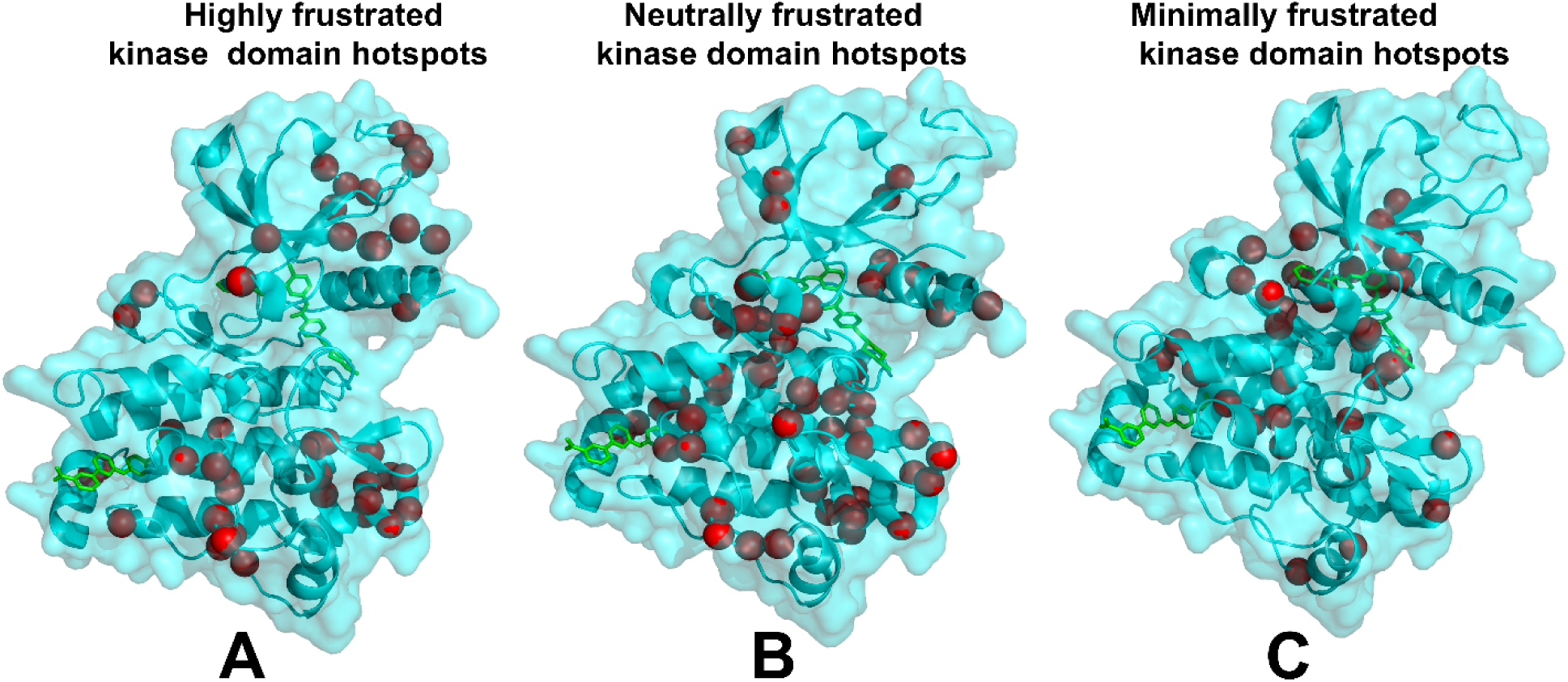
Structure-based analysis and mapping of local frustration patterns in the dataset of protein kinase complexes including both orthosteric (type I/II) and allosteric (type III/IV) complexes. Structural mapping of highly frustrated kinase domain residue hotspots (the top 10% of highly frustrated sites) (A), neutrally frustrated kinase residue hotspots (the top 10% of neutrally frustrated sites) (B) and minimally frustrated kinase residue hotspots (the top 10% of minimally frustrated sites) (C). The kinase domain is shown in light, pink-colored ribbons and the high frustration hotspots are shown in red spheres. Structural mapping of local frustration hotspots is projected onto the inactive ABL kinase domain complex with the type II Imatinib and allosteric inhibitor GNF-2 (pdb id 3K5V). Imatinib and GNF-2 inhibitors are shown in sticks. GNF-2 binds to the myristoyl pocket in the C-lobe.

Hence, our results indicate that highly frustrated kinase domain positions may have evolved strategically to create small nucleation sites promoting large conformational changes. The emergence of local frustration in these regions could also indicate the “initiation cracking points” of the inactive kinase form, ultimately facilitating global conformational transitions and shifting a dynamic equilibrium between kinase states towards the active form.^65^ Local cracking model in protein kinases is often referred to formation of unfavorable residue-residue interactions that can be compensated by increased local entropy in intermediate states during large conformational changes between inactive and active states.^65,66^

Strikingly, our analysis reveals that the global frustration profiles of orthosteric and allosteric kinase complexes are indistinguishable in their overall architecture. This similarity indicates that the global energetic framework of the kinase catalytic domain is largely conserved, irrespective of the ligand binding mode. Within this shared framework, neutral frustration dominates the catalytic domain, with relative residue densities peaking sharply in the range of 0.55–0.75 (Figure 7). The neutrally frustrated positions occupy dense areas across the entire domain, particularly along the central spine connecting the N– and C-terminal lobes, enabling a balance of conformational plasticity and mutational tolerance (Figure 8B). The prevalence of neutral frustration may reflect a fundamental biophysical principle: kinases must maintain sufficient conformational flexibility to toggle between active and inactive states yet avoid destabilizing energy barriers that would impede functional transitions. Neutral frustration provides precisely this balance permitting functional structural rearrangements without affecting the fold stability and protecting structural core of the kinase catalytic domain.

Minimally frustrated residues—indicative of evolutionarily optimized, energetically stable interactions—also constitute a substantial fraction of the domain, with relative densities spanning 0.1–0.6 before rapidly decaying to baseline (Figure). These residues are enriched in structurally conserved elements such as the β-sheet core, catalytic binding site, catalytic loop, and regulatory spine, consistent with their role in maintaining global fold integrity (Figure 8C). These observations align closely with general principles of protein energy landscapes, which consistently report that ∼50–60% of local contacts are neutrally frustrated, ∼30% minimally frustrated, and only ∼10% highly frustrated across diverse protein folds. The kinase catalytic domain—despite its intricate regulatory mechanisms—adheres to this universal pattern, underscoring that functional complexity does not require energetic exceptionality, but rather strategic placement of frustration within a predominantly neutral scaffold.

However, beneath this global uniformity lies a critical local divergence at functional binding sites. The comparative analysis of local frustration across orthosteric and allosteric kinase sites reveal a striking dichotomy – while the global energy landscape of the kinase catalytic domain remains broadly conserved, the local frustration profiles at functional binding sites exhibit profound differences that map directly onto evolutionary constraint and predictive model performance (Figure 9). The distributions of frustration density for residues within orthosteric binding sites (Type I/II inhibitors targeting the ATP cleft). show a pronounced shift toward minimal frustration (Figure 9).

**Figure 9.**
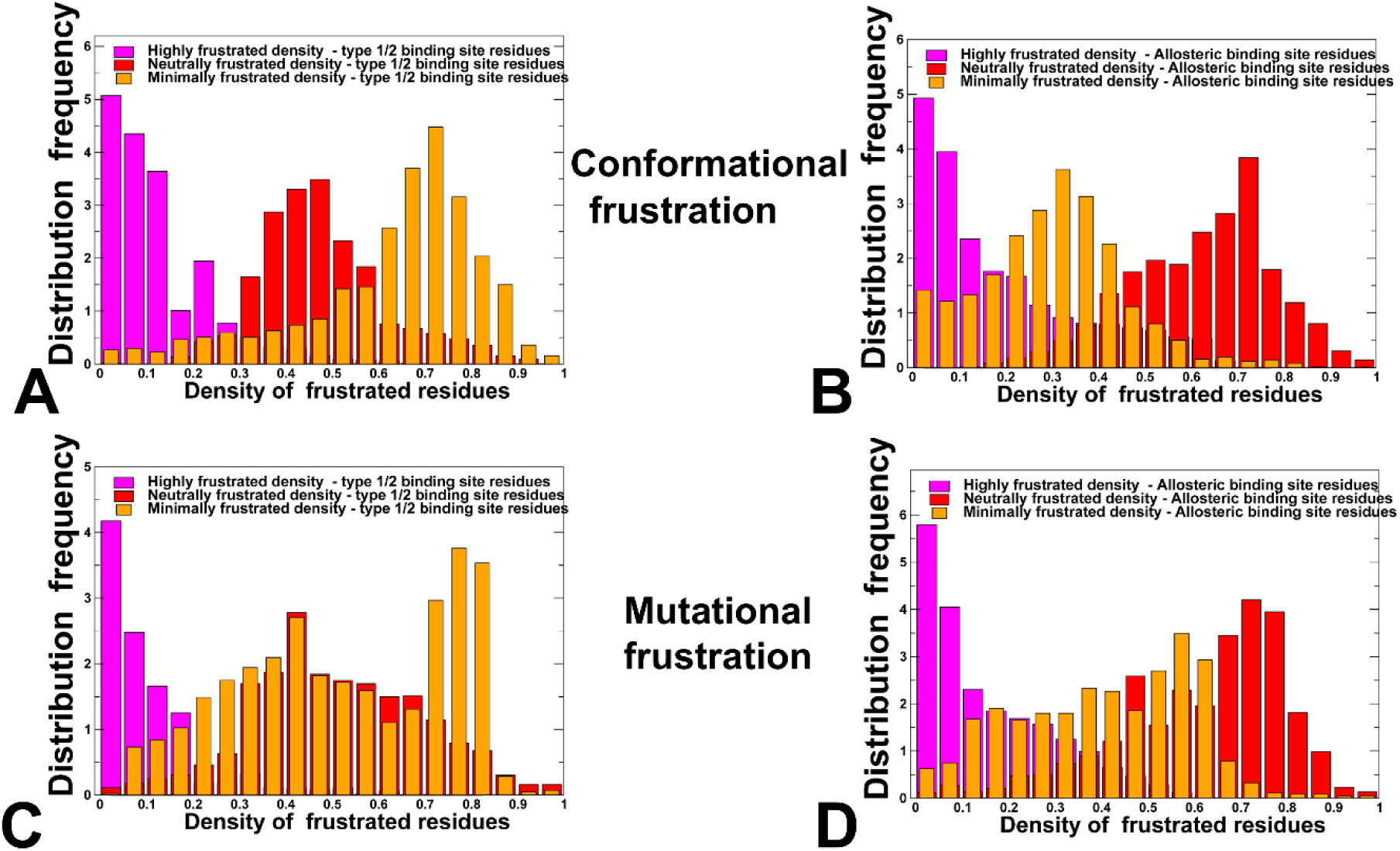
The distributions of conformational and mutational frustration of the type 1/II orthosteric and type III/IV allosteric binding site residues in the protein kinase complexes. The relative global densities of conformational frustration for orthosteric binding site residues in the protein kinase complexes (A) allosteric binding site residues in the protein kinase complexes (B). The relative global densities of mutational frustration for orthosteric binding site residues in the kinase complexes (C) allosteric binding site residues in the kinase complexes (D). The highly frustrated density of kinase domain residues is shown in magenta-colored filled bars, neutrally frustrated density is in red-colored bars and minimally frustrated density is in orange-colored bars.

For conformational frustration, the distribution peaks sharply in the high-density region (>0.7-0.75), indicating that native interactions at these positions are strongly stabilized compared to decoys generated by local structural perturbations (Figure 9A). This reflects the more rigid, well-defined geometry required for ATP binding and catalysis. Even more dramatically, the mutational frustration distribution is heavily skewed toward minimal frustration (Figure 9C). The native contacts in these regions are energetically optimized for catalytic function as mutational substitutions or geometric perturbations become largely destabilizing. This signature reflects intense selection on catalytic residues, which must preserve precise geometry for ATP coordination and phosphor-transfer. The resulting sequence conservation generates a strong unambiguous evolutionary signal for PLMs, trained on deep coevolutionary patterns, resulting in robust detection of orthosteric binding sites in protein kinases with high fidelity. Structure-based methods such as P2Rank further benefit from the geometric regularity and surface prominence of these sites.

In stark contrast, allosteric sites, whether proximal (Type III) or distal (Type IV)are characterized by predominant neutral local frustration and an elevated, though still minor, population of highly frustrated residues (Figure 9B,D). Conformational frustration is dominated by neutral frustration (red bars, peak ∼0.65–0.75), there is a noticeable increase in the population of highly frustrated residues (magenta bars) compared to orthosteric sites (Figure 9C). For mutational frustration, we similarly observed a pronounced shift toward neutral mutational frustration suggesting that amino acids involved in the formation of diverse allosteric pockets can undergo functional conformational rearrangements and be evolutionarily permissive, allowing for sequence drift while preserving its capacity for conformational switching or ligand-induced remodeling (Figure 9D). Crucially, there is also a small but significant tail of highly frustrated residues – positions where even the native amino acid is energetically unfavorable, suggesting these are “hotspots” of regulatory tension may be involved in conformational switching or ligand-induced stabilization.

This profile indicates that residue identities at allosteric positions are evolutionarily permissive: many substitutions are energetically neutral, enabling sequence drift while preserving conformational plasticity. Functionally, this permissiveness allows kinases to evolve new regulatory inputs—new ligand sensitivities or protein–protein interfaces—without compromising core catalysis. We argue that this adaptability renders allosteric sites more difficult for robust predictions as PLM may often view diversity allosteric sites as functional noise as surface patches lacking conserved motifs, exhibiting high sequence variability, and indistinguishable from the vast sea of neutral, non-functional residues.

### 3.4. Class Imbalance Alone Cannot Explain Markedly Reduced Performance of PLM on Allosteric Sites

The collapse of precision (AUPR ≈ 0.06) on allosteric kinase complexes, even as ranking ability remains moderate (AUROC ≈ 0.70) cannot be attributed solely to extreme class imbalance (<3% binding residues). While sparsity undoubtedly exacerbates the problem, three lines of evidence point to a potentially more fundamental cause – the intrinsic biophysical and evolutionary design of allosteric sites renders them inherently ambiguous to sequence-based models, regardless of data quantity or balance. First, our PLM was trained on the large, diverse, LIGYSIS dataset, which includes tens of thousands of binding residues across orthosteric and allosteric sites from hundreds of protein families. Despite this broad exposure, the model fails specifically on non-ATP pockets, indicating that the issue is not merely insufficient allosteric examples, but the absence of a consistent, learnable signal. Scaling the training data (more sequences, more structures) reinforces the model ability to find consensus signals (conservation/minimal frustration). Because allosteric sites are diverse, structurally heterogeneous, and lack a universal sequence motif, more data could potentially increase the “noise” for family-specific allosteric sites rather than lead to measurable improvements. Second, class imbalance cannot explain performance disparities within the allosteric spectrum. Type III pockets (prevalence = 4.8%, AUPR = 0.29) are as abundant as Type I.5 sites (prevalence = 6.0%, AUPR = 0.76), yet their precision is less than half. This gap arises not from data scarcity, but from differences in mutational constraint: Type III sites already exhibit neutral mutational frustration and structural heterogeneity, which erode coevolutionary signals even when examples are plentiful.

Third, the lack of robust PLM predictions of the allosteric sites is not necessarily due to an inability to model neutrally frustrated regions. In fact, PLMs excel at predicting the structure of the kinase catalytic domain across nearly all regions including large swaths of the N– and C-lobes that are dominated by neutral conformational frustration, just like allosteric sites. This success stems from the fact that the kinase fold itself is evolutionarily overrepresented in training data as its core β-sheet, αC-helix, and regulatory spines are highly conserved across >500 human kinases. Thus, even neutrally frustrated residues within this shared, recurrent structural scaffold benefit from implicit evolutionary priors that stabilize predictions. In contrast, allosteric sites, whether proximal (Type III) or distal ( type IV ALLO), are not part of this conserved architectural core. They often reside in flexible loops (e.g., A-loop, P+1 motif), dynamic hinges or lineage-specific pockets (e.g., ABL myristoyl site) that are underrepresented, structurally divergent, and functionally non-conserved across the human kinome. As a result, while the PLM can accurately model the structural fold of a kinase domain, including its neutrally frustrated functional regions, the model can encounter significant difficulties in inferring the functional relevance of a surface patch that lacks coevolutionary coupling, structural recurrence, or catalytic constraint. In other words, neutral frustration is only “interpretable” when embedded in a conserved structural context while in isolated structural context, it may appear as noise. This distinction clarifies why AUROC remains moderate (0.70–0.89) for allosteric sites. Indeed, the model can weakly rank them above random surface residues but cannot assign high confidence because they lack the minimal mutational frustration signature that generates unambiguous evolutionary signals. Following these arguments, it is possible that even in a perfectly balanced dataset where allosteric and orthosteric sites each constituted 50% of residues, the PLM may still assign low scores to allosteric residues, as their sequence context is statistically indistinguishable from the vast sea of non-functional, neutrally frustrated surface residues.

Our results lead to a conjecture that evolution may have intentionally masked allosteric sites to preserve regulatory fidelity where allosteric sites may be under active selection for conformational and sequence ambiguity—a strategy we term “evolutionary concealment.” While orthosteric sites are locked into highly conserved, minimally frustrated sequences to ensure catalytic fidelity across phylogeny, allosteric sites appear to be deliberately diversified to avoid spurious activation or inhibition by endogenous or environmental molecules. In this interpretation, the “invisibility” of allosteric sites to coevolutionary signals may be an intrinsic functional feature. Why would natural selection favor such concealment? Allosteric sites are inherently context-dependent and combinatorial to enable nuanced responses to specific signals—phosphorylation states, binding partners, metabolic cues, or localized drug concentrations. We argue that by maintaining broad neutral frustration and allowing many amino acid substitutions and dynamic changes without energetic penalty, evolution can ensure that allosteric sites remain plastic, tunable, and lineage-specific. Our frustration analysis reveals that these sites may function as energetically primed “cryptic rheostats,” embedded within a shared conformational scaffold but partly shielded from evolutionary exposure. To detect them, models may need to incorporate biophysical signals such as frustration landscapes that encode the energetic grammar of functional plasticity.

### 3.5. ABL Kinase as a Canonical Playground for Decoding Blind Spots in Allosteric Binding Site Prediction

To ground our global frustration analysis in atomic detail and validate its interpretive power of PLM and P2Rank predictions of binding, we performed residue-level profiling across a structurally diverse ensemble of ABL kinase complexes—spanning active, intermediate, and fully autoinhibited states. This system serves as an ideal paradigm because ABL regulatory mechanism is exquisitely characterized, its catalytic domain is representative of the tyrosine kinase fold, and it hosts both orthosteric inhibitors (e.g., Dasatinib, Axitinib) and well-defined allosteric modulators (e.g., GNF-2, DPH, Asciminib) that stabilize distinct conformational states.

We present a detailed residue-level analysis of high, neutral, and minimally frustrated positions across distinct ABL kinase complexes: (a) the active and inactive catalytic domain conformations formed by type I kinase inhibitors Dasatinib (pdb id 2GQG)^67^ and Axitinib (pdb id 4WA9)^68^; (b) the inactive ABL structure bound with type II inhibitor Imatinib and allosteric inhibitor GNF-2 (pdb id 3K5V)^69^, inactive ABL complex with type II Imatinib and allosteric activator DPH (pdb id 3PYY)^70^ and the autoinhibitory ABL regulatory complex with type II inhibitor Nilotinib and allosteric inhibitor Asciminib (pdb id 5MO4).^71,72^

This analysis serves as atomic-scale validation of our central thesis: orthosteric and allosteric sites are divergently optimized by evolution for stability versus plasticity, and this divergence is encoded in their local frustration signatures, which in turn may dictate the success or failure of PLM predictors. We explore several critical, interconnected purposes that directly anchor our biophysical interpretation to the main computational findings of this study. First, ABL provides a structurally and functionally canonical model system for dissecting the energetic signatures of orthosteric versus allosteric regulation. The ABL kinase catalytic domain is a canonical representative of the tyrosine kinase family and shares the conserved bi-lobal architecture common to protein kinases (Figure 10). Conformational dynamics and transitions between the inactive and active ABL kinase states are orchestrated by the conserved 361-HRD-363 motif in the catalytic loop and the 381-DFG-383 motif that are coupled with the regulatory αC-helix (residues 279-291) to form regulatory spine (R-spine) (L301,361,F382,D421) and catalytic spine (C-spine) (A269, L323, C369, L370, F317) networks (Figure 10). Dynamics and stabilization of these networks serve as a strong indicator of protein kinase activation. The well-characterized conformational landscape of ABL spanning active, partially inactive, and fully autoinhibited states enables direct mapping of frustration patterns to ligand type, functional outcome (activation vs. inhibition), and binding site class (orthosteric vs. allosteric). By anchoring global trends to a well-understood kinase paradigm, we provide mechanistic credibility and structural interpretability of PLM and P2Rank prediction results.

**Figure 10.**
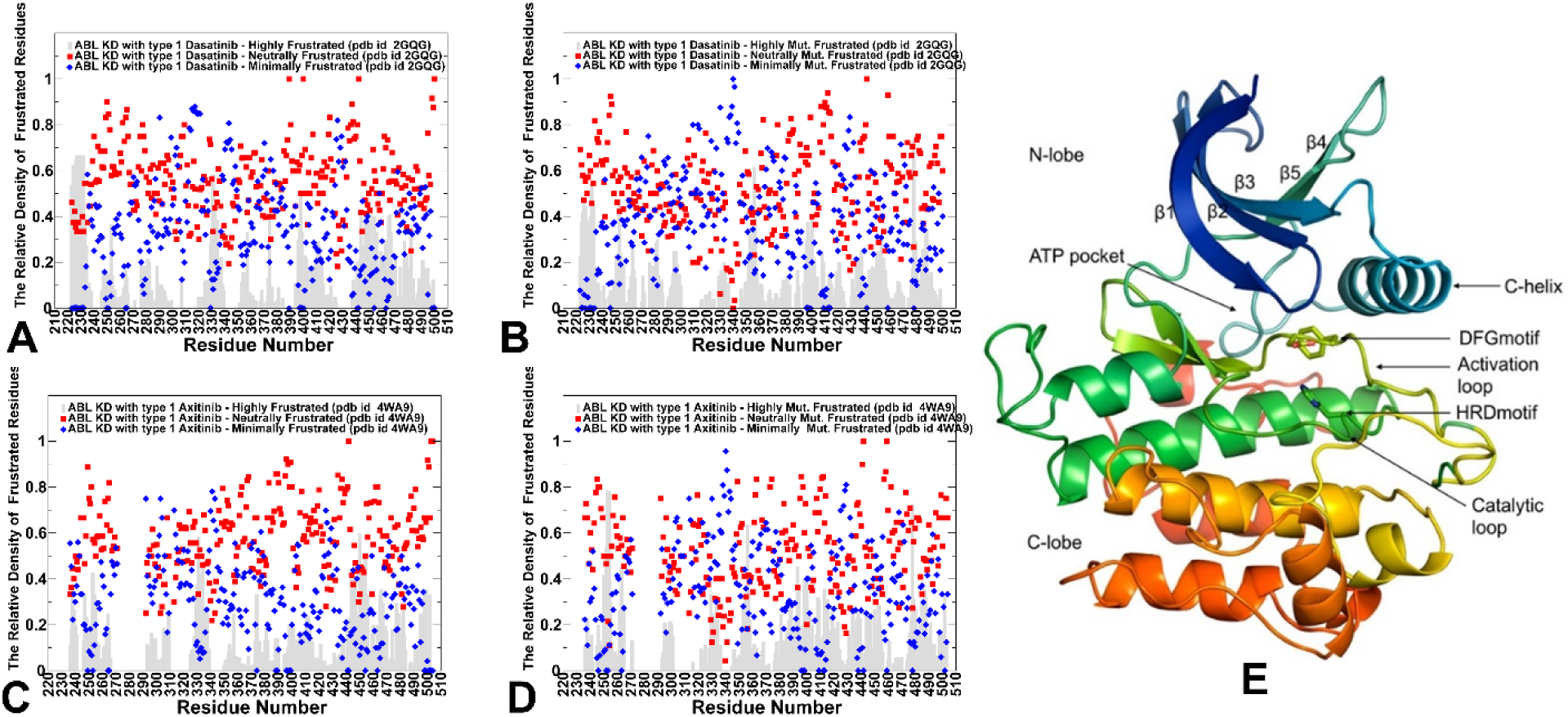
The residue-based distributions of conformational. (A,C) and mutational frustration (B,D) for ABL complexes with type I inhibitors Dasatinib (A,B) and Axitinib (C,D). The relative densities of high frustration are shown in grey background bars, the relative densities of neutral frustration in red squares and minimal frustration are in magenta diamonds. (E) The generic structure of the active form of the kinase catalytic domain (in orange ribbons) with key functional regions annotated and pointed by arrows including ATP binding site, C-helix, DFG motif, HRD motif, A-loop, and catalytic loop.

We first characterized conformational and mutational frustration profiles of ABL kinase residues in complexes with type I orthosteric ATP-competitive inhibitors binding active state (Figure 10).

By analyzing the Dasatinib complex, we capture the archetypal orthosteric site: evolutionarily conserved and geometrically optimized (Figure 10A,B). The conformational frustration profile for this complex revealed the overall dominance of neutrally frustrated residues over the kinase domain with a notable concentration of minimally frustrating cluster of residues near the catalytic site (residues 315-321) that constitute the core of inhibitor-interacting sites (Figure 10A). Of some notice, the presence of several clusters of high frustration density (residues 320-340 in C-lobe, A-loop residues 395-420, the P+1 loop with WTAPE motif (residues 400-410) and the αG-helix (residues 440-455) which serves as a scaffold for A-loop (Figure 10A). The WTAPE motif is embedded in the middle segment of the activation loop and is Positioned just upstream of the P+1 substrate-binding loop and downstream of the DFG and HRD motifs. In the active (DFG-in) state, the WTAPE motif adopts a β-turn-like structure that helps anchor the A-loop in an open, extended conformation, facilitating substrate access. Although mutational frustration residue profile for type I kinase complex with Dasatinib (Figure 10B) is similar to the corresponding conformational frustration profile (Figure 10A), we observed the increased concentration of mutational minimally frustrated residues.

A similar conformational and mutational frustration profiles were seen for another prototypical kinase domain complex with the type I kinase inhibitor Axitinib (Figure 10C,D). Dasatinib binds deep in the ATP cleft, forming key H-bonds and hydrophobic contacts with T315, F317, M318, Y320, G321, L370, M290, L299 (Figure 10A,B,E). Axitinib forms four hydrogen bonds with ABL interacting with T315, K271 and Y253, F382 residues (Figure 10C,D.E). The examination of the frustration profiles for both type I inhibitors targeting orthosteric ATP binding site reveals significant presence of dominant minimally frustrated positions near ATP region (residues 310-330) (Figure 10A,C). We noticed stronger presence of mutational minimal frustration in these positions, indicating considerable evolutionary conservation in the orthosteric binding site region (Figure 10B,D).

The emergence of moderate but noticeable high frustration density peals reflects the intrinsic nature of the kinase landscape that has encoded specific kinase domain regions with “frustration potential” that can be modulated through minimal-neutral-high zones to trigger switching between alternative functional states. The key observation from the frustration analysis of type I inhibitor-kinase complexes is a clear preference of minimally frustrated positions in the orthosteric binding site where both Dasatinib and Axitinib are surrounded by active site residues (Figure 10).

Structural mapping of corresponding high, neutral, and minimal frustrated positions on the crystal structure of ABL complex with Dasatinib highlights the key finding, whereby the largest density corresponds to neutrally frustrated positions and minimally frustrated residues with the latter concentrated near the ATP binding site (Figure 11). The smaller and more localized clusters of potentially highly frustrated positions are sprinkled through C-lobe, overlapping with locations of various allosteric pockets, and populating the αG-helix (residues 440-455) which serves as a scaffold for triggering functional changes of the A-loop and activation transitions.

**Figure 11.**
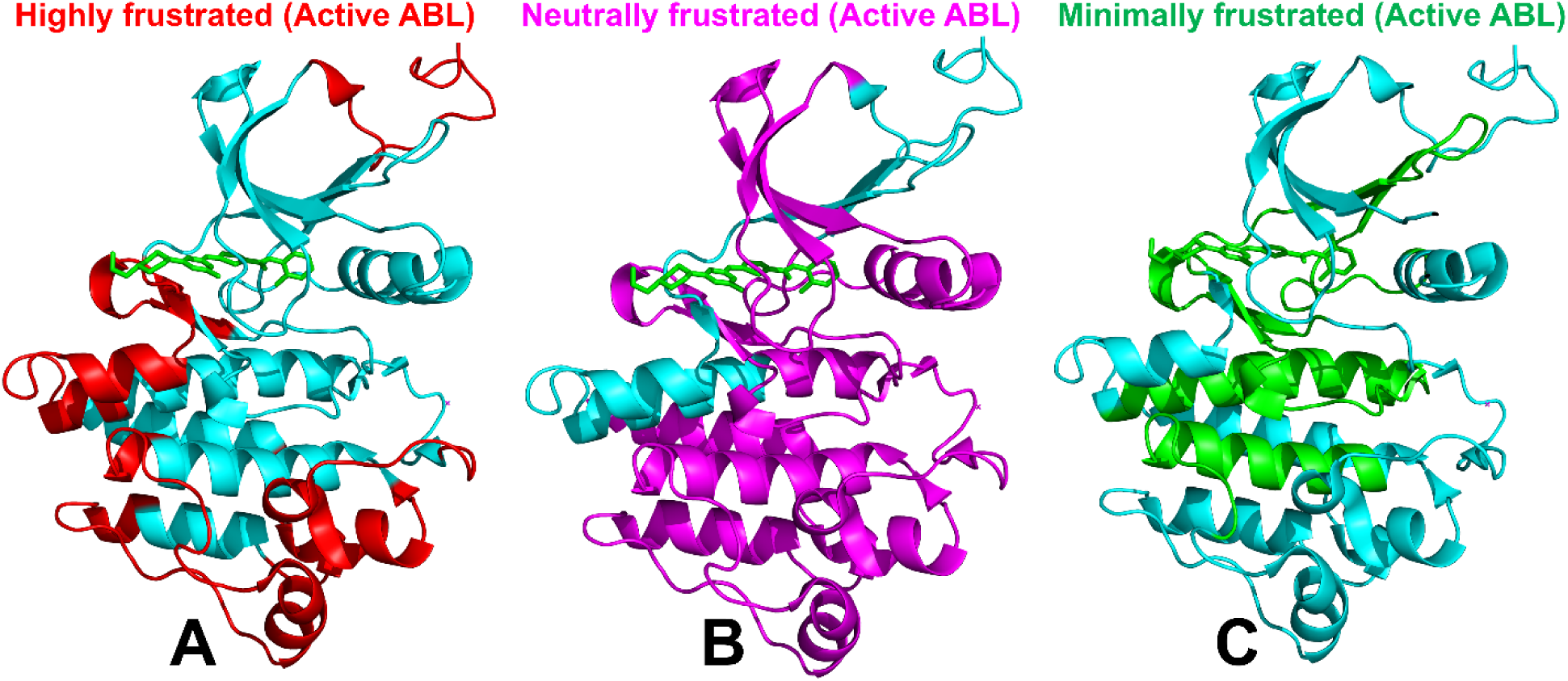
Structural mapping of local frustration densities in the ABL structure with type I Dasatinib and Axitinib inhibitors. The structural map of highly frustrated ABL kinase domain residue hotspots (the top 10% of highly frustrated sites) (A), neutrally frustrated kinase residue hotspots (the top 10% of neutrally frustrated sites) (B) and minimally frustrated kinase residue hotspots (the top 10% of minimally frustrated sites) (C). The kinase domain is shown in light, pink-colored ribbons and the high frustration density is shown in red on panel A, neutral frustration density in magenta on panel B, and minimal frustration in green on panel C. A prototypical type I Dasatinib inhibitor is shown in sticks.

To sum up, in the type I-bound active conformation the ATP-binding cleft is enriched in minimally frustrated residues forming a contiguous, energetically optimized hotspot that directly contacts the inhibitor. Mutational frustration is even more pronounced in minimally frustrated positions in this region than conformational frustration, reflecting strong selection as nearly any substitution at these positions is energetically destabilizing. This creates unambiguous, high-fidelity evolutionary signal that PLM can readily detect explaining their high precision on Type I orthosteric sites.

Of notable importance is the residue profiling of conformational and mutational frustration in the inactive form of the Abl kinase domain, bound in a complex with both the ATP-competitive inhibitor imatinib and the allosteric inhibitor GNF-2 (Figure 12A,D), ABL complex with Imatinib and allosteric activator DPH (pdb id 3PYY) (Figure 12B,E) and ABL complex the ABL regulatory complex with Nilotinib and allosteric inhibitor Asciminib (pdb id 5MO4) (Figure 12C,F), The presence of both inhibitors in the structure helps to lock the protein into this down-regulated, inactive state. The conformational frustration profiles (Figure 12A-C) and mutational frustration profiles (Figure 12D-F) for these three inactive ABL complexes reveal a similar unifying theme whereby conformational profiles are dominated by neutrally frustrated residues while mutational frustration distributions also feature notably significant populations of minimally frustrated positions. Despite the overall prevalence or neutral frustration, there are also notable clusters of highly frustrated contacts that include A-loop (384–410) residues folding inward and making extensive contacts with the αG-helix (425–445). The P+1 motif (∼404–412) resides at the interface between these two elements. Together, they form a contiguous frustrated region characterized by non-native residue–residue contacts and energetic strain. This tripartite assembly is critical for allosteric control: its formation locks ABL in an inactive state, and its disruption (e.g., by myristoyl-pocket inhibitors like GNF-2) can either stabilize or destabilize autoinhibition depending on context. Hence, our results indicated that the ABL kinase regions undergoing large structural changes during inactive-active transitions could be enriched in clusters of highly frustrated residues. These findings are consistent with recent studies showing that identification of conserved highly frustrated contacts can determine residues involved in conformational transitions. The emergence of highly frustrated clusters in the αC-helix, A-loop and C-lobe regions could present the “initiation cracking points” that could perturb the inactive state and promote dynamic functional transitions to the catalytically competent active kinase form.

**Figure 12.**
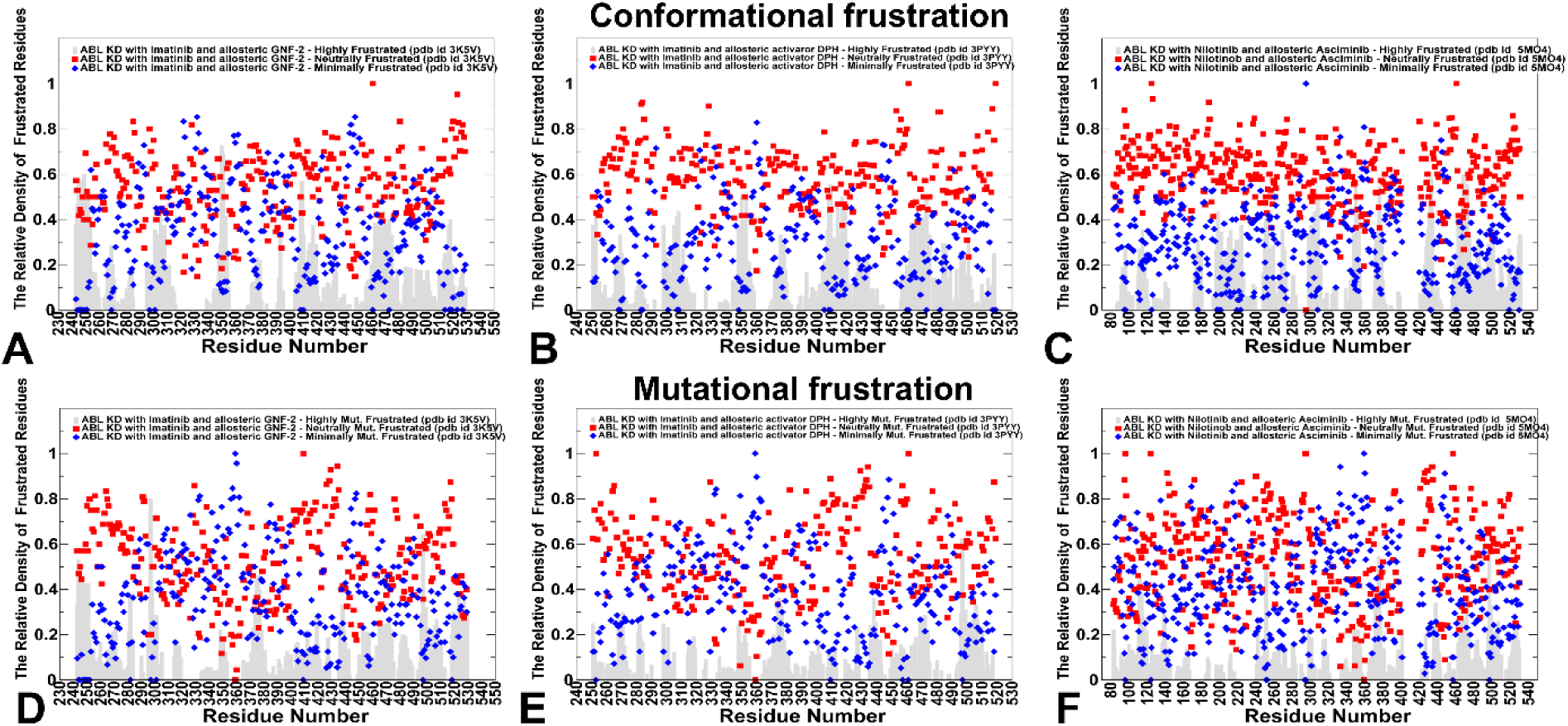
The residue-based distributions of conformational. (A-C) and mutational frustration (D-F) for ABL complexes with type II inhibitors and allosteric inhibitors/activators. (A) Conformational frustration profiles for the inactive ABL structure bound with type II inhibitor Imatinib and allosteric inhibitor GNF-2 (pdb id 3K5V) (panel A), inactive ABL complex with type II Imatinib and allosteric activator DPH (pdb id 3PYY) (panel B) and the ABL complex with type II inhibitor Nilotinib and allosteric inhibitor Asciminib (pdb id 5MO4) (panel C). Mutational frustration residue profiles for the ABL kinase domain in the respective complexes are shown on panels (D-F). The relative densities of high frustration are shown in grey background bars, the relative densities of neutral frustration in red squares and minimal frustration are in magenta diamonds.

Central to our analysis is the general structural conservation of regions enriched by high, neutral, and minimally frustrated residues. Structural mapping indicates that neutrally and highly frustrated positions may be enriched near the ABL allosteric myristoyl-pocket as well as near other potential allosteric pocket locations present in other kinases (Figure 13A,B).

**Figure 13.**
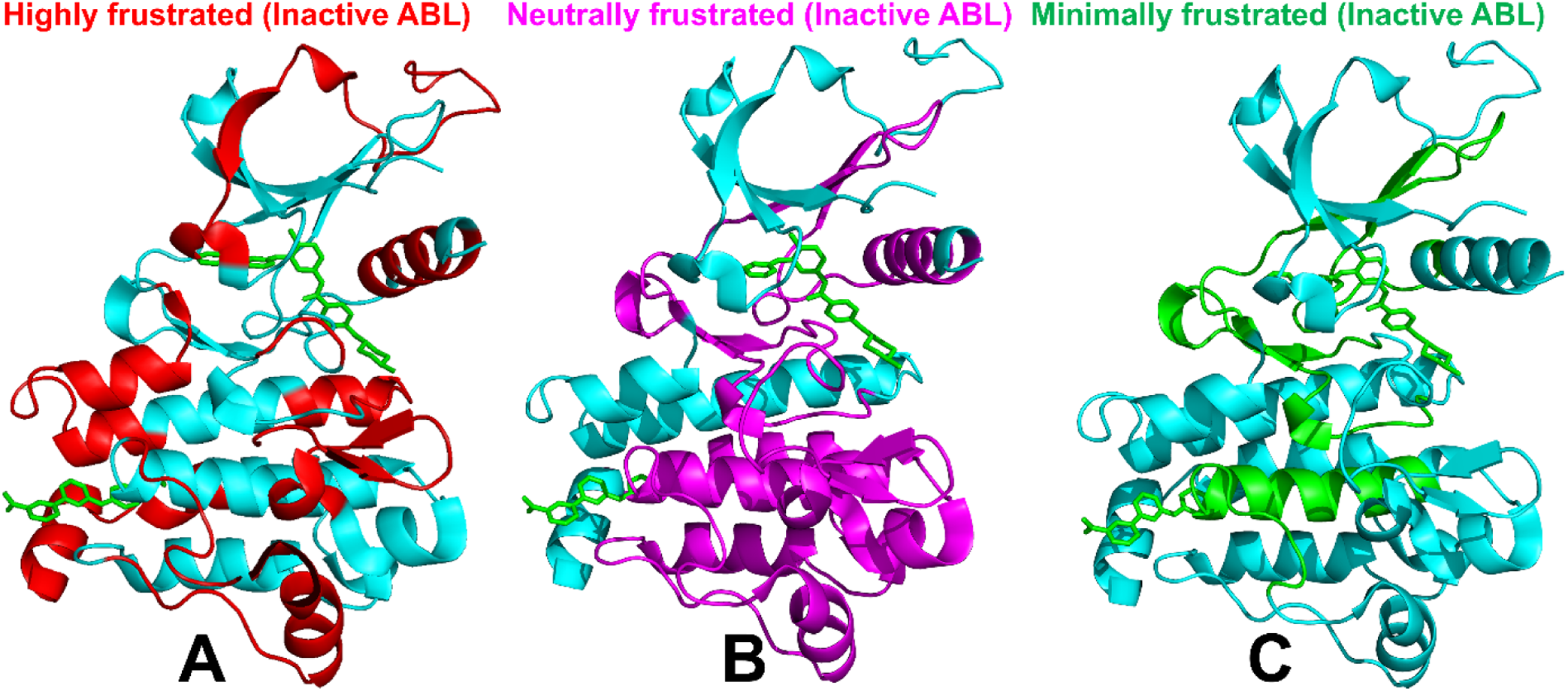
Structural mapping of local frustration densities in the inactive ABL kinase domain bound to the ATP-competitive type II inhibitor Imatinib and the allosteric inhibitor GNF-2 (pdb id 3K5V). The structural map of highly frustrated ABL kinase domain residue hotspots (the top 10% of highly frustrated sites) (A), neutrally frustrated kinase residue hotspots (the top 10% of neutrally frustrated sites) (B) and minimally frustrated kinase residue hotspots (the top 10% of minimally frustrated sites) (C). The kinase domain is shown in light, pink-colored ribbons and the high frustration density is shown in red on panel A, neutral frustration density in magenta on panel B, and minimal frustration in green on panel C. A prototypical type II Imatinib inhibitor and allosteric inhibitor GNF-2 are shown in sticks.

At the same time, the minimally frustrated density is featured prominently for ATP binding sites residue in all complexes. Notably, mutational frustration profile reveals even more pronounced preference for minimal frustration in the orthosteric binding site residues (Figure 13C). These ABL-specific findings provide atomic-scale validation of our central thesis: orthosteric sites are minimally frustrated and evolutionarily constrained, while allosteric sites are neutrally frustrated and evolutionarily permissive. We argue that the divergence in frustration profiles stems from fundamentally different evolutionary pressures. Orthosteric sites are under strong negative selection, leading to minimal mutational frustration and high sequence conservation. On the other hand, allosteric sites may be evolutionary designed to be context-dependent and modifiable. The prediction dependence on coevolutionary informational signals explains the lack of predicted power for diverse and conformationally dynamic allosteric sites with often context-dependent mix of neutrally and highly frustrated positions that are not readily discernible by learning of coevolutionary signals in protein families. While PLMs excel at detecting evolutionary constraint, this signal may be intentionally minimized at allosteric positions that can manipulate the extent of structural and mutational variations to optimize for function.

### 3.6. Analyzing Predictions of Protein Kinase Binding Sites through the Lens of Energy Landscape-Encoded Frustration as Explainable AI Framework

Accurate identification of functional allosteric pockets remains a central challenge in structure-based drug discovery. In this study, we benchmarked a fine-tuned protein language model (PLM) against the structure-based predictor P2Rank across a rigorously curated dataset of human kinase–ligand complexes. The results reveal a systematic divergence: PLMs achieve high precision on orthosteric ATP-binding sites (AUPR > 0.7) but perform poorly on allosteric sites (AUPR < 0.1), despite retaining moderate ranking ability (AUROC ≈ 0.70). This pattern holds consistently across Type I, I.5, and II inhibitors—agents that target the conserved ATP cleft—and breaks down for Type III and non-ATP allosteric modulators, which bind structurally heterogeneous, evolutionarily labile pockets. Crucially, this performance gap is not attributable to class imbalance alone—though allosteric sites constitute fewer than 3% of kinase residues—but reflects a deeper biophysical reality. To interpret this divergence, we integrated large-scale local frustration analysis, a framework rooted in energy landscape theory that quantifies how native residue–residue interactions compare energetically to decoy ensembles generated by mutational or conformational perturbations. This analysis reveals that predictive success maps directly onto the evolutionary and energetic signatures of functional sites.

Both orthosteric and allosteric kinase complexes share a conserved global frustration landscape: neutral frustration dominates (∼55–75% of residues), with substantial contributions from minimally frustrated contacts and only a sparse fraction of highly frustrated residues. This architecture reflects an evolutionarily optimized balance—sufficient conformational plasticity to enable transitions between active and inactive states, yet enough stability to avoid non-productive unfolding. However, at the local level, binding sites exhibit starkly divergent frustration profiles.

Orthosteric sites—encompassing the ATP cleft and its extensions—are characterized by pronounced minimal mutational frustration. Substitutions at these positions are energetically destabilizing, reflecting strong purifying selection on residues essential for catalysis and ATP coordination. This constraint produces a clear, high-fidelity evolutionary signal in sequence space—one that PLMs, trained on deep multiple sequence alignments, are exquisitely tuned to detect. The resulting conservation enables reliable residue-level prediction even from sequence alone. In contrast, allosteric sites display predominant neutral mutational frustration, with a modest but significant enrichment of highly frustrated residues. Neutral mutational frustration indicates that many amino acid substitutions are energetically tolerated—evidence of relaxed evolutionary constraint. Functionally, this permissiveness allows kinases to evolve context-dependent regulatory mechanisms without compromising core catalysis. Yet this same adaptability renders allosteric sites invisible to sequence-based models. To a PLM, these regions appear as functional noise: surface patches lacking conserved motifs, exhibiting high sequence variability, and indistinguishable from the vast sea of neutral, non-functional residues.

This dichotomy reframes the challenge of allosteric site prediction: it is not a failure of AI, but a mismatch between model assumptions and biological design. PLMs excel at detecting evolutionary constraint—a feature intentionally minimized at allosteric sites, which operate as tunable rheostats rather than rigid switches. Their functional relevance is encoded not in sequence conservation, but in energetic plasticity: the capacity to adopt alternative conformations, form cryptic pockets, and respond to ligand-induced remodeling.

Our findings have several key implications. First, frustration analysis provides an essential interpretability layer for AI-driven binding site prediction. Rather than treating model output as a black-box score, one can contextualize predictions through physical principles: high-confidence PLM predictions in minimally frustrated regions likely reflect true orthosteric sites; low-confidence predictions in neutrally frustrated regions may signal cryptic allosteric hotspots. This diagnostic capability could improve target prioritization and reduce false leads in early-stage screening. Second, the results argue for a paradigm adjustment in binding site prediction: from purely data-driven models to biophysics-integrated AI. We propose that energetic frustration metrics should not be only post-hoc interpretability tools—but integral features in next-generation predictive models. By embedding frustration indices (configurational and mutational) as priors or constraints within PLM fine-tuning or graph neural network architectures, we can explicitly teach models to “look for” regions of neutral frustration that are primed for conformational switching. Such models would no longer rely solely on evolutionary echoes but on physical principles of functional plasticity. It also reframes the goal of binding site prediction: not just to *locate* pockets, but to classify their regulatory potential based on biophysical signatures. Allosteric drugs offer superior selectivity and reduced toxicity, yet their discovery remains largely serendipitous due to the invisibility of their targets to conventional computational screens. Our framework also offers a diagnostic: when a PLM confidently predicts a binding site in a minimally frustrated region, it likely reflects a conserved, druggable orthosteric cleft. When it fails in a neutrally frustrated region, that failure itself may be a signal—an indicator of a cryptic, evolutionarily labile, but potentially high-value allosteric site.

Future predictors could embed frustration metrics as inductive biases—e.g., by weighting attention entropy in transformer layers based on mutational frustration, or by training graph neural networks on frustration-guided conformational ensembles that explicitly model pocket-opening transitions. Such approaches would move beyond pattern recognition toward mechanistic prediction. From a drug discovery perspective, this analysis may provide interesting implications.

This study demonstrates that the performance of AI models on binding site prediction is not determined by algorithmic sophistication alone, but by alignment with the biophysical logic of protein function. By grounding predictions in energy landscape theory, we transform model limitations into mechanistic insights and compass for rational allosteric drug discovery.

## 4. Conclusions

This study presents a comprehensive evaluation of residue-level binding site prediction in protein kinases using both a fine-tuned protein language model (PLM) and the structure-based method P2Rank. Across a rigorous dataset spanning multiple kinase inhibitor classes, both approaches achieved high accuracy for orthosteric sites—particularly Type I, I.5, and II ATP-competitive pockets—demonstrating the robustness of current AI strategies when strong evolutionary or structural signals are present. In contrast, performance on allosteric sites (Type III and non-ATP pockets) was markedly reduced, with precision collapsing to near-baseline levels despite moderate ranking capability. This divergence persisted even after accounting for class imbalance, dataset leakage, and conformational diversity, underscoring a fundamental limitation in how sequence– and structure-based models encode functional information.

To interpret this performance gap, we integrated large-scale local frustration analysis—a physics-based framework rooted in energy landscape theory—that quantifies the energetic stability of residue–residue interactions under mutational and conformational perturbations. Critically, our results show that the global frustration landscape of the kinase catalytic domain is conserved across ligand types, underscoring a shared architecture of conformational plasticity. This analysis revealed a striking biophysical dichotomy: while the global frustration landscape of the kinase catalytic domain is conserved, orthosteric and allosteric sites differ fundamentally in their mutational constraints. Orthosteric pockets are enriched in minimally frustrated residues, reflecting strong purifying selection and yielding the clear evolutionary signal that PLMs leverage. Allosteric sites, by contrast, are dominated by neutral mutational frustration, indicating evolutionary permissiveness that enables conformational plasticity at the cost of sequence conservation. This energetic ambiguity renders them largely invisible to PLMs and only partially accessible to static structure-based methods like P2Rank.

Together, these findings establish local frustration as a robust interpretability layer that bridges AI predictions and biophysical reality. Rather than viewing low precision on allosteric sites as model failure, our analysis reframes the predictive performance of PLM and P2Rank tools as a reflection of functional design that can be rationalized and improved through lens of energetic and geometric frustration of biological systems. The convergence of AI benchmarking and energy landscape analysis thus provides a principled framework for diagnosing prediction outcomes, prioritizing cryptic sites, and guiding the development of next-generation models that embed physical priors—such as frustration-guided attention or ensemble-aware architectures—to capture the dynamic, context-dependent nature of regulatory binding.

To conclude, this study suggests that reliable detection of allosteric pockets could require not only more scaling data or model parameters; it demands an integration of statistical learning with biophysical insight. It provides a principled framework to diagnose model limitations, prioritize biologically relevant targets, and ultimately bridge the gap between sequence, structure, and function in the pursuit of next-generation allosteric therapeutics.

## Data Availability Statement

Data is fully contained within the article and Supplementary Materials. Crystal structures were obtained and downloaded from the Protein Data Bank (http://www.rcsb.org). The rendering of protein structures was done with UCSF ChimeraX package (https://www.rbvi.ucsf.edu/chimerax/) and Pymol (https://pymol.org/2/).

The code and dataset are available in GitHub repository: https://github.com/skrhakv/LBS-pLM. The fine-tuned PLM model can be downloaded from this data storage: https://owncloud.cesnet.cz/index.php/s/8RSJqt60D2uWJNa

Inference was performed using custom Python scripts that wrap the fine-tuned ESM2 checkpoint and implement residue extraction, tokenization, batching, and model evaluation. All scripts required to reproduce the PLM predictions reported here are publicly available in GitHub repository (https://github.com/kiarka7/plm-vs-p2rank-kinases). The fine-tuned model checkpoint can be provided upon request for reproducibility.

The data, including the trained models, can be downloaded at https://cunicz-my.sharepoint.com/:u:/g/personal/40001559_cuni_cz/Ebf0PKmYSD9NvLGdH5pl-e0BuNUM8hJFTfM0zMm4-tac9g. The detailed description of the data organization can be found in Supplementary Materials.

## Author Contributions

All authors have read and agreed to the published version of the manuscript.

