## Supplementary Text for "Protein Language Models and Structure-Based Machine Learning for Prediction of Allosteric Binding Sites in Protein Kinases: An Explainable AI Framework Grounded in Energy Landscape-Encoded Frustration"

The data, including the trained models, can be downloaded at https://cunicz-my.sharepoint.com/:u:/g/personal/40001559_cuni_cz/Ebf0PKmYSD9NvLGdH5pl-e0BuNUM8hJFTfM0zMm4-tac9g

The data include

2) Data files (per kinase family) (in shared folder):

in each family directory (ALLO, Type I, Type I.5, Type II, Type III):

-01.chain_gold_labels_CLEAN.json – ligand-proximal ground truth (4 Å heavy-atom criterion) for clean dataset

-02.chain_p2rank_labels.json – P2Rank CSV outputs and original predictions (p2rank_predictions folder)

-03.plm_predictions.json – raw PLM residue-level probabilities

-03.chain_plm_labels.json - labels

-04.eval_all_in_one_CLEAN.json – all metrics for all methods on the CLEAN subset

3) Global metadata (in shared folder)

-clean dataset in .xlsx – full list of chains filtered out by UniProt overlap

-kinase_uniprot_map_typeX.tsv – retained chains after leakage filtering

-00.clean_summary.csv – retention statistics per family

4) Scripts (in shared folder)

All Python scripts with brief comments are uploaded in this folder. All scripts and software developed and used in this study are fully and freely available at GitHub: https://github.com/kiarka7/plm-vs-p2rank-kinases

5) Software

The fine-tuned PLM can be provided upon request. P2Rank is available in https://github.com/rdk/p2rank.

Binding-site residues for each kinase structure

All binding-site labels used for both PLM and P2Rank evaluation are already included in the dataset you received.

They are in the following JSON files (one set per kinase family):

-01.chain_gold_labels_CLEAN.json — residues derived directly from PDB structures

-02.chain_p2rank_labels.json — P2Rank pocket predictions

-03.chain_plm_labels.json — PLM predictions at the fixed threshold

*03.plm_predictions.json — raw PLM probabilities and list of residues with probability ≥ 0.75

Each of these files contains, for every (pdb_id, chain_id):

-sequence_residue_numbers — residue indices used for alignment

-labels — 0/1 vector (GOLD or P2Rank)

-pred_labels_fixed — 0/1 vector for PLM predictions (in 03.chain_plm_labels.json)

*additionally, in 03.plm_predictions.json one would also find the following information : residues_predicted — residue numbers where $p \geq 0.75$ and probabilities — per-residue PLM raw scores.

The files 02.chain_p2rank_labels.json and 03.chain_plm_labels.json contain all original (unfiltered) structures.

For the CLEAN dataset used in the analysis, only structures and chains present in 01.chain_gold_labels_CLEAN.json were selected and evaluated. Therefore, to extract binding-site residues corresponding strictly to the CLEAN subset, P2Rank and PLM files must be filtered by the (pdb_id, chain_id) pairs listed in 01.chain_gold_labels_CLEAN.json.
